# Genetic code expansion enables programmable covalent protein design

**DOI:** 10.64898/2026.05.15.725538

**Authors:** Helena de Puig, Erkin Kuru, Michaël Moret, Allison Flores, Suneesh Karunakaran, Dinara Sayfullina, Subhrajit Rout, Carmen Escobedo-Lucea, James J. Collins, George M. Church

## Abstract

Covalent chemistry has transformed small-molecule drug discovery, yet analogous strategies for proteins remain largely inaccessible because covalent warheads cannot be readily integrated into biologics. Conventional genetic code expansion requires engineering a dedicated aminoacyl-tRNA synthetase for each new amino acid, rendering broad warhead screening impractical. Here we introduce AminoX, a platform that bypasses this limitation through direct tRNA acylation, enabling site-specific incorporation of chemically diverse non-standard amino acids (nsAAs), including covalent warhead nsAAs compatible with scalable biologic manufacturing and multifunctional nsAAs. Using a pooled mRNA display workflow, we screened more than 2,000 warhead-position combinations in machine learning-designed *de novo* miniproteins targeting CTLA-4, enabling parallel interrogation of covalent chemistry, linker geometry, and incorporation site. We confirmed covalent engagement on cells together with enhanced functional blockade. Finally, we demonstrate multifunctional nsAAs that combine covalent warheads with fluorogenic reporters for real-time detection of target engagement, as well as dual nsAA incorporation for macrocyclization and fluorescent imaging of covalent binding on cell surfaces. By uniting synthetic biology, chemical biology, generative protein design, and high-throughput functional selection, AminoX compresses covalent protein engineering timelines by orders of magnitude, accelerating the development of next-generation therapeutics, biosensors, and chemical probes.

## Introduction

Machine learning-based protein design has transformed our ability to generate miniproteins with tailored specificities and nanomolar affinities against therapeutic targets ^1–6^. Their designability, small size, stability, and resistance to proteases make them attractive candidates for next-generation biologics. However, miniproteins are restricted to the 20 canonical amino acids, limiting their chemical functionality to what evolution has provided—a stark contrast to peptides, where non-standard amino acids (nsAAs) are routinely incorporated through solid-phase synthesis ^7^.

Genetic code expansion (GCE) offers a route to transcend the genetic code’s alphabet by enabling site-specific incorporation of nsAAs bearing functionalities absent from the canonical repertoire such as electrophilic warheads for covalent target engagement, fluorophores for visualization, click handles for bioconjugation, and crosslinkers for macrocyclization ^8–14^. However, traditional GCE applied to proteins depends notably on engineered aminoacyl-tRNA synthetases (aaRSs), each accommodating only a narrow set of substrates. Engineering a new aaRS requires extensive work with no guarantee of success, making systematic exploration of chemical space impractical. While flexizyme-based approaches have demonstrated aaRS-free incorporation ^15–17^, they have been largely confined to peptides and linear scaffolds. Translating this capability into an efficient high-throughput discovery platform that systematically screens diverse chemistries, positions, and scaffolds in parallel remains an open challenge.

Separately, mRNA display has emerged as a powerful platform for selecting functional molecules from large libraries by covalently linking each encoded variant to its own mRNA, enabling direct sequencing-based readout of enriched sequences ^18,19^. The technology has been used extensively to discover macrocyclic peptide binders against therapeutic targets ^17,20^, and more recently to screen covalent peptides ^21^. However, both applications have been largely confined to peptide scaffolds, leaving larger, folded proteins (the scaffolds of interest for most biologics) underexplored. Combining direct tRNA acylation with pooled mRNA display would in principle allow parallel screening of diverse nsAA chemistries, positions, and scaffolds in folded proteins, but this integration has not previously been demonstrated. A recent yeast-display platform has enabled selection of fast-acting covalent protein drugs by combining display libraries with post-translational chemoselective modification, yielding covalent nanobodies, miniproteins, and cytokines with rapid crosslinking kinetics ^22^. This approach screens large libraries against one warhead chemistry at a time, with reactive groups installed post-translationally on cysteine residues. What has not been addressed is the complementary problem: screening hundreds of combinations of warhead chemistry and incorporation sites in one pooled experiment to find which drive covalent engagement.

Here, we introduce AminoX, an aaRS-free platform that integrates direct chemical acylation of tRNAs ^23–25^, miniaturized cell-free translation, machine learning-guided protein design ^1^, and pooled mRNA display. Through efficient direct chemical acylation, AminoX bypasses aaRS engineering entirely, enabling high-throughput incorporation of chemically diverse nsAAs into diverse scaffolds such as nanobodies and *de novo*–designed miniproteins to unlock functionalities previously inaccessible to folded protein scaffolds. Freedom from synthetase constraints also enables nsAAs to be designed from commodity precursors like lysine, linking discovery to scalable manufacturing of both the nsAA and the final biologic.

To demonstrate our platform’s capabilities, we focused primarily on covalent target engagement, which has had transformative impact on small-molecule drug discovery. Drugs such as ibrutinib, osimertinib, and sotorasib achieve durable clinical responses through irreversible binding and sustained target occupancy ^26,27^. Pioneering work by Wang and colleagues has extended this concept to proteins through proximity-enabled reactive therapeutics (PERx), demonstrating that genetically encoded electrophilic nsAAs can form selective covalent bonds with target nucleophiles in antibodies and nanobodies ^11,28,29^. Proximity-induced covalent nsAAs remain inert until positioned near a target nucleophile, where they form selective irreversible bonds that enhance binding durability, selectivity, and pharmacodynamic control ^30^. However, these efforts have relied on individually engineered synthetases for each nsAA-containing warhead, limiting systematic optimization of warhead chemistry and linker geometry. Covalent engagement of *de novo*–designed miniproteins has been limited to individual warhead–position combinations installed on pre-existing binders ^31,32^; systematic exploration across warhead chemistries and incorporation sites has not been reported.

We designed nine chemically distinct covalent warhead amino acids spanning a range of linker geometries and electrophile chemistries. Using a pooled mRNA display workflow, we screened 612 warhead × position combinations in a known miniprotein binder against CTLA-4 and identified dozens high-confidence covalent hits at a target with no previously characterized covalent engagement sites. We further extended this workflow to *de novo* miniprotein design with machine learning: we designed CTLA-4 binders with BindCraft ^1^, quantified binding in our cell-free pipeline, and screened 1,458 position × warhead combinations across three lead scaffolds, identifying covalent variants with reduced dissociation rates and enhanced functional blockade in cellular assays.

Beyond covalency, we demonstrated the generality of the platform through dual nsAA incorporation, installing both a covalent warhead and a fluorescent reporter into a single protein at two distinct positions for real-time tracking of target engagement on live cells. We further demonstrate intramolecular macrocyclization via click chemistry between two incorporated nsAAs. We also introduce fluorogenic warheads that combine covalent reactivity with turn-on fluorescence into a single nsAA design, enabling optical readout of covalent bond formation. Together, these expansions position AminoX as a foundation for engineering multifunctional biologics of any size, with capabilities that approach those of small molecules.

## Results

### AminoX enables covalent warheads with tunable geometry

AminoX builds on three complementary technologies: chemical acylation of tRNA, cell-free protein synthesis, and orthogonal codon decoding. Any non-standard amino acid (nsAA) is first conjugated to a 5′-phospho-2′-deoxyribocytidylylriboadenosine (pdCpA) dinucleotide and ligated onto a truncated orthogonal tRNA, typically an amber-suppressor, yielding an aminoacylated tRNA_CUA_ without the need for an evolved synthetase. The charged tRNA is then used in a cell-free protein synthesis (CFPS) system, typically PURExpress ΔRF1, where it decodes an in-frame amber (UAG) codon introduced at any defined position in the target protein ^23^. Because the system works directly in the reaction tube, proteins can be expressed, tested, and iterated in microliter volumes within hours.

To establish that covalent warheads can be incorporated through this workflow, we reproduced two previously characterized covalent binders. The first, LCB3 (PDB ID: 7JZM) ^31,32^, is a miniprotein that covalently binds the SARS-CoV-2 spike RBD (Fig. 1A,B). The second, KN035 (PDB ID: 5JDS) ^33,34^, is a single-domain antibody that engages PD-L1 (Fig. 1C,D). Both proteins engage their target covalently using fluorosulfate-L-tyrosine (FSY), a well-characterized proximity-enabled reactive (PERx) electrophilic nsAA that reacts with nearby histidine, tyrosine, and lysine residues ^28^.

**Fig. 1.**
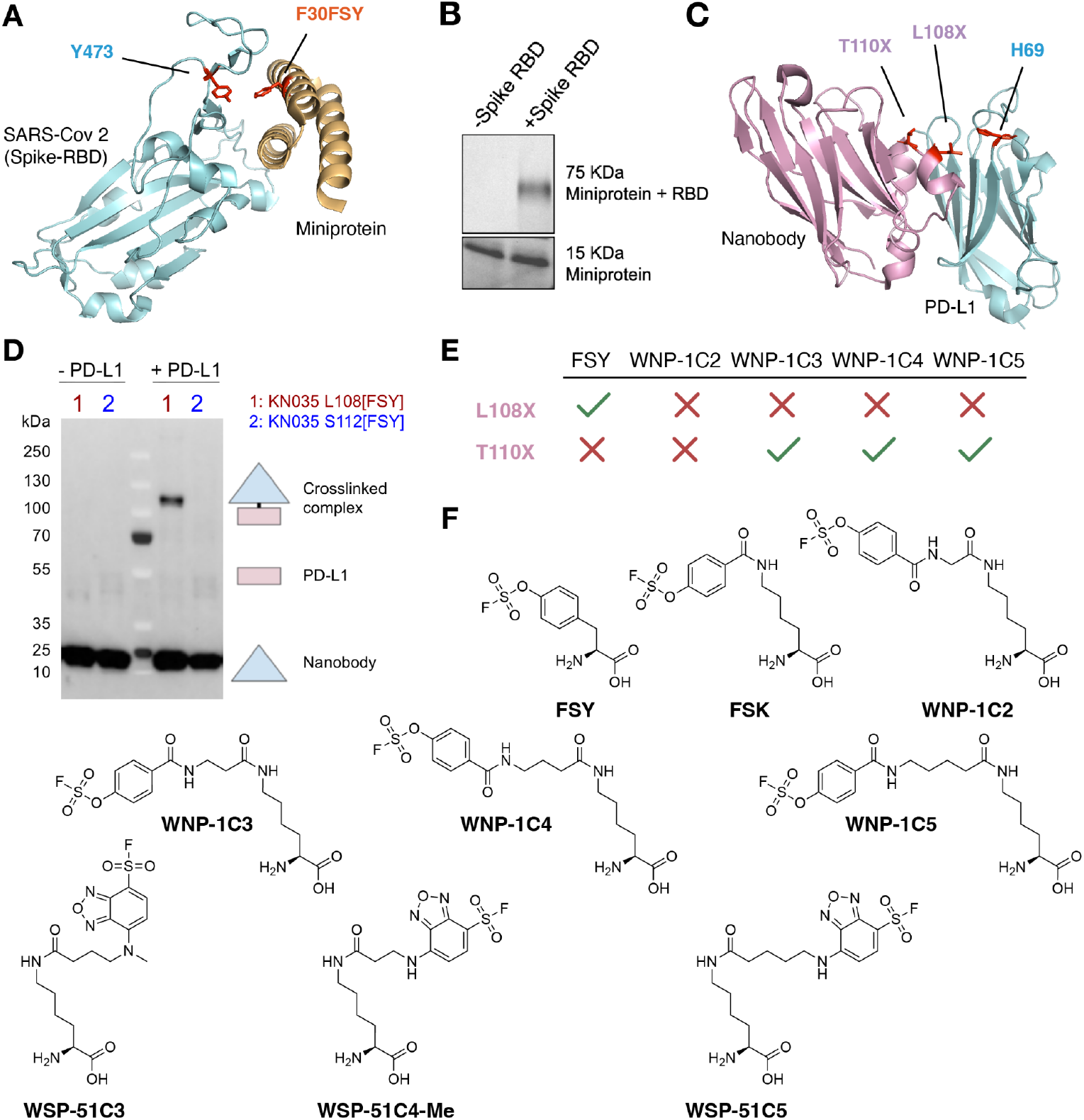
Rapid generation of covalent miniprotein and nanobody binders against SARS-CoV-2 RBD and PD-L1. **A**. Crystal structure of the SARS-CoV-2 spike receptor-binding domain (RBD) in complex with a miniprotein binder (PDB ID: 7JZM) that was subsequently converted into a covalent binder. The position of the covalent warhead and the targeted residue on RBD are highlighted in red. **B**. Gel electrophoresis of covalent miniprotein showing adduct formation in the presence but not absence of spike RBD. **C**. Crystal structure of PD-L1 in complex with a nanobody (KN035, PDB ID: 5JDS) that was subsequently converted into a covalent binder. **D**. Western blot of nanobodies with FSY incorporated at positions 108 or 112, showing covalent adduct formation in presence of PD-L1 at position 108. **E**. Position-dependent covalent engagement to PD-L1 of the KN035 nanobody with FSY and novel FSK derivatives with diverse linker lengths. **F**. Chemical structure of FSY, FSK/1C1, novel FSK derivatives (WNP-1C2, WNP-1C3, WNP-1C4, WNP-1C5), and novel covalent fluorogenic reporters (WSP-51C3, WSP-51C4-Me, WSP-51C5).

We first confirmed FSY incorporation by expressing a model peptide and verifying the expected mass by HPLC-MS (Supplementary Fig. 3). Next, we tested FSY reactivity against RBD. Gel electrophoresis of heat-denatured protein complexes showed that LCB3 containing FSY at position 30 (LCB3 F30[FSY]) formed a higher-molecular weight band, corresponding to a covalently linked LCB3-RBD complex only observed in the presence of RBD. This confirmed target-specific covalent bond formation (Fig. 1B). For the single-domain antibodies (KN035), we started by showing that binding was preserved in CFPS without purification (2 nM purified, 9 nM not purified, Supplementary Fig. 2), allowing us to accelerate the time from DNA to binding characterization with biolayer Interferometry (BLI). We then incubated purified PD-L1 with unpurified CFPS reactions expressing KN035 108[FSY] and KN035 112[FSY]. Western blot (WB) analysis showed that KN035 108[FSY] formed a covalent complex with PD-L1, with the observed molecular weight matching the sum of PD-L1 and KN035 (Fig. 1D, Supplementary Fig. 1). KN035 112[FSY] did not form a covalent complex (Fig. 1D), consistent with prior observations ^33^. Together, these results establish that covalent warheads are compatible with our direct-acylation workflow and that CFPS-derived proteins retain the covalent reactivity required for target engagement.

Because AminoX decouples warhead development from synthetase evolution, new nsAAs can be designed, synthesized, and encoded within days rather than the months-to-years typically required to evolve a dedicated synthetase. We leveraged this to generate a series of fluorosulfate amino acids with different linker lengths (Fig. 1F), turning warhead geometry into a tunable design variable rather than a fixed constraint. After confirming site-specific incorporation of WNP-1C2, WNP-1C3, WNP-1C4, and WNP-1C5 by LC-MS (Supplementary Fig. 3), each differing by a single methylene unit, we applied this to a structurally demanding covalent site: position 110 of the PD-L1-binding nanobody (14.2 Å from PD-L1 H69, Fig. 1C), which sits beyond the theoretical reach of FSY and FSK (∼9 Å).

When incorporated at position 110, these extended nsAAs successfully enabled covalent crosslinking to PD-L1 (Fig. 1E), as demonstrated by the appearance of a higher molecular weight band on WB corresponding to the nanobody–PD-L1 adduct (Supplementary Fig. 4). Systematic tuning of linker length therefore opens up covalent sites that are geometrically inaccessible with fixed-geometry warheads.

### High-throughput discovery of covalent hotspots by pooled mRNA display

Having shown that single warhead–position combinations can be engineered at known covalent sites, we next asked whether the platform could discover covalent hotspots (binder positions whose incorporated warheads covalently engage a target residue) at scale, across hundreds of nsAA–position combinations in parallel.

To that end, we selected cytotoxic T-lymphocyte–associated protein 4 (CTLA-4), and focused on the *de novo* designed miniprotein 8GAB, which binds CTLA-4 with ∼0.1 nM affinity and has a solved co-crystal structure (PDB ID: 8GAB) ^35^. Because 8GAB has no previously identified covalent engagement sites, any hits would represent *de novo* covalent positions on an existing high-affinity binder.

To systematically probe covalent engineering opportunities across the 8GAB–CTLA-4 interface, we developed an mRNA display workflow that enables single-step covalent selection in folded proteins. Prior mRNA display approaches to covalent discovery have been limited in two ways. Early efforts avoided selecting reactive warheads directly by using two-step workarounds (selecting a non-reactive bait first, then chemically converting it to a warhead post-selection) ^21^; more recent single-step approaches have been restricted to cyclic peptide scaffolds ^36–38^.

Our initial attempts confirmed the challenge. False positives appeared, which we attributed to two factors: high-temperature steps inherent to typical mRNA display workflows, and unreacted warhead carried into the selection that produced spurious crosslinks. We confirmed the temperature effect by incubating covalent nanobodies (KN035 L108[FSY] & KN035 L112[FSY]) with PD-L1 at different temperatures: higher temperatures increased reactivity but decreased specificity, with non-specific crosslinking to cell-free components observed at elevated temperatures (Supplementary Fig. 5). We therefore designed a protocol that combines low-temperature conditions throughout with a quenching step using excess free tyrosine and histidine to block unreacted warheads to bias selection toward true warhead–target adducts.

Next, we introduced amber (UAG) codons at 68 positions proximal to the 8GAB binding interface and translated each library with nine different nsAAs: five lysine-derived WNP warheads (FSK/1C1, WNP-1C2, WNP-1C3, WNP-1C4, WNP-1C5), three novel fluorogenic NBD-SF (7-nitrobenzo-2-oxa-1,3-diazole sulfonyl fluoride) warheads (WSP-51C3, WSP-51C4-Me, WSP-51C5) verified by LC-MS for site-specific incorporation (Supplementary Fig. 18), as well as FSY as a benchmark (Fig. 1F). The platform also supports barcoded reverse transcription primers that encode nsAA identity into the cDNA, allowing libraries from different warheads to be pooled and competed in a single selection, then deconvoluted by sequencing (Fig. 2A). This generated a library of 612 potential covalent protein variants (68 positions × nine nsAAs). Using our quenching protocol, after incubation with CTLA-4 and stringent washes, we re-translated the enriched DNA pool with each nsAA individually and confirmed single-round selection of nsAA-specific covalent binders, validating the approach (Supplementary Fig. 6).

**Fig. 2.**
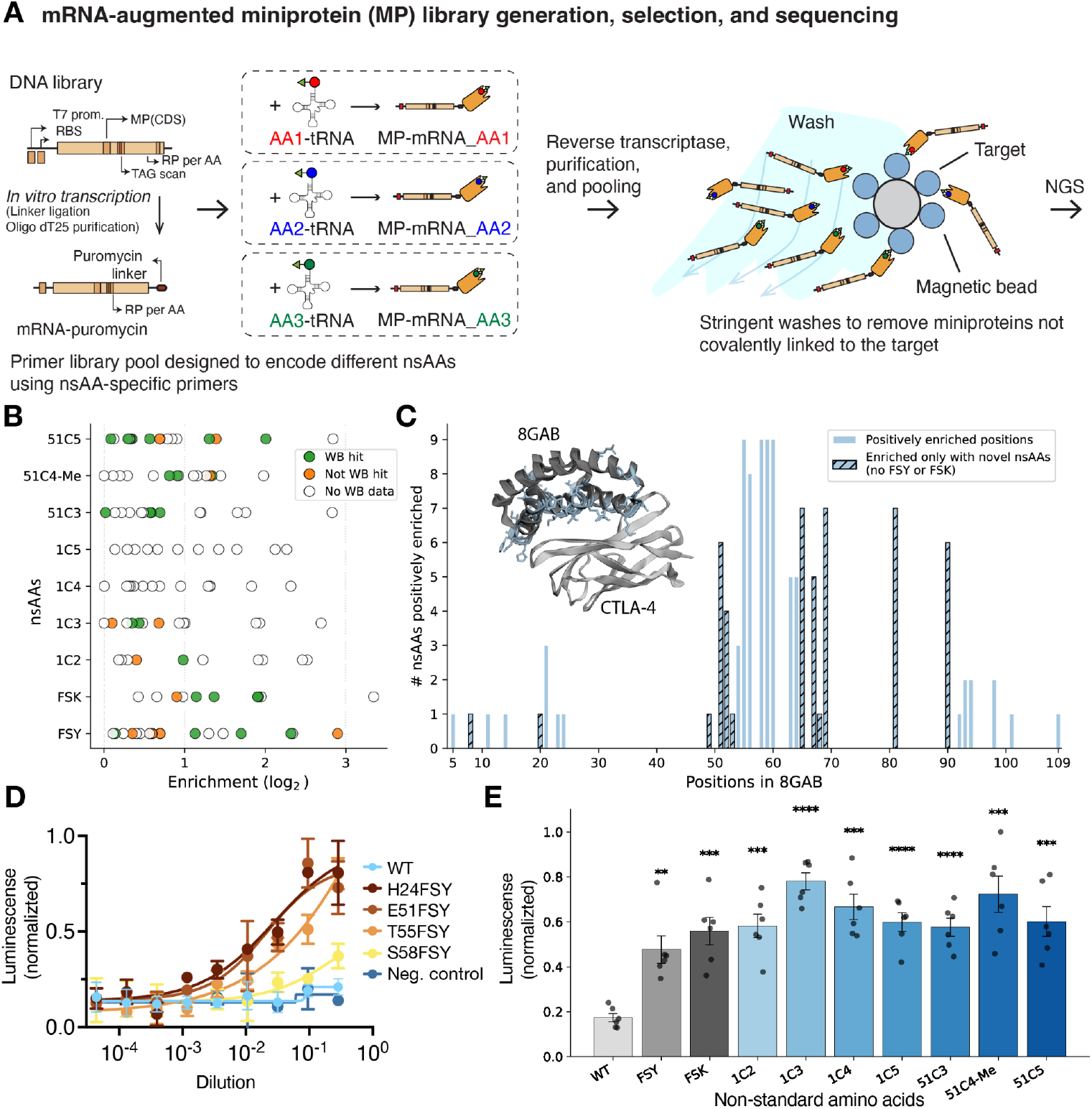
Pooled mRNA display identifies covalent hotspots in 8GAB and reveals functional enhancement of CTLA-4 blockade. **A**. Scheme for mRNA display library preparation. TAG scan libraries are translated with distinct nsAA-tRNA_CUA_ pairs and covalently linked to their encoding mRNA. Reverse transcription with nsAA-specific barcoded primers generates mRNA/cDNA–protein fusions for pooled selection. In total, 612 variants (68 positions × nine nsAAs) were incubated with CTLA-4, quenched with excess free tyrosine and histidine, then subjected to stringent washes including boiling. **B**. Positively enriched variants by nsAA. Green dots also showed positive hits in western blot (WB), orange dots did not appear as hits in WB, white circles were not tested in WB. **C**. Positively enriched positions on 8GAB (light blue). Hatched bars are positioned only enriched with novel amino acids designed in this work. **D**. Functional validation with CTLA-4 MOA Promega assay of various covalent FSY-miniprotein variants at different dilutions (8GAB H24[FSY], 8GAB E51[FSY], 8GAB T55[FSY], 8GAB S58[FSY]) showing enhanced potency compared to wild-type (8GAB WT, wild type, light blue). **E**. CTLA-4 MOA Promega assay of 8GAB H24X at 10X dilution modified with nine different nonstandard amino acids. 10µl of nsAA containing H24X 8GAB were expressed by CFPS for 4h at 30C followed by incubation for 6h at 37C. Increased luminescence indicates stronger blockade. Bars show the mean of six replicates ± SEM, with individual replicates overlaid as dots. Statistical significance versus WT was assessed by Welch’s two-sided t-test: p ≥ 0.05; *, p < 0.05; **, p < 0.01; ***, p < 0.001; ****, p < 1×10^-4^.

Sequencing revealed broad enrichment across the binding interface (Fig. 2C). Of the 612 nsAA × position pairs in the scanned library, 121 (∼20%) showed positive log_2_ enrichment after selection, distributed across 32 of the 68 scanned positions. Four sites (55, 58, 59, and 60) were positively enriched with all nine nsAAs tested, marking a continuous belt of permissive residues at the binding interface. A second hot region spanned positions 63–69 (notably 65 and 69, hit by seven of nine nsAAs each), and additional clusters appeared at 51–52 and 90–94. Importantly, twelve positions were positively enriched only with the novel nsAAs, with no enrichment for the established warheads FSY or FSK (positions 8, 20, 49, 51, 52, 53, 65, 67, 68, 69, 81, 90). These sites suggest geometries that the canonical warheads cannot reach but the novel chemistries can.

To compare the sequencing readout against a biochemical standard, we cross-validated 159 variants across the scanned library by WB (Supplementary Figures 7-12); 77 (∼48%) registered as covalent hits. Restricting to positively enriched variants (log_2_ > 0; Fig. 2B), 25 of 36 WB-tested variants (69%) were independently confirmed as covalent hits. The cut therefore enriches for true positives, but it is not a complete filter. Conversely, among confirmed hits, NGS enrichment provides a more fine-grained basis for ranking candidates than WB band intensity, which is difficult to compare quantitatively across blots and between gels; the most highly enriched variants are therefore prime priorities for downstream characterization. We also verified that our mRNA display workflow extends beyond miniproteins to nanobodies, single-chain variable fragments (scFvs), and heavy-chain antibodies. They all successfully displayed as mRNA–protein fusions against PD-L1 and CTLA-4 (Supplementary Fig. 13).

Functional evaluation using a cell-based mechanism-of-action assay (CTLA-4 MOA Promega assay) revealed that covalent proteins were stronger inhibitors than the non-covalent variants. Proteins expressed in 10 µL PURExpress reactions were directly diluted into cell media without purification and assayed. While wild-type unpurified, non-covalent 8GAB showed low activity in disrupting CTLA-4–dependent cell-cell interactions, several covalent variants demonstrated robust functional enhancement (Fig. 2D, E). Importantly, both covalent reactivity and functional efficacy varied depending on incorporation site and covalent amino acid structures, highlighting the value of our systematic screening approach and modular nsAA design capability (Fig. 2D, E).

### End-to-end covalent miniprotein design from target to binder

Having converted a pre-existing high-affinity binder into covalent variants, we next tested whether the platform could generate covalent binders starting from a target but without a validated binder scaffold. This is the more general and more demanding case: most therapeutic targets lack a starting binder like 8GAB, and both binding specificity and covalent reactivity must be generated rather than tuned (Fig. 3A).

**Fig. 3.**
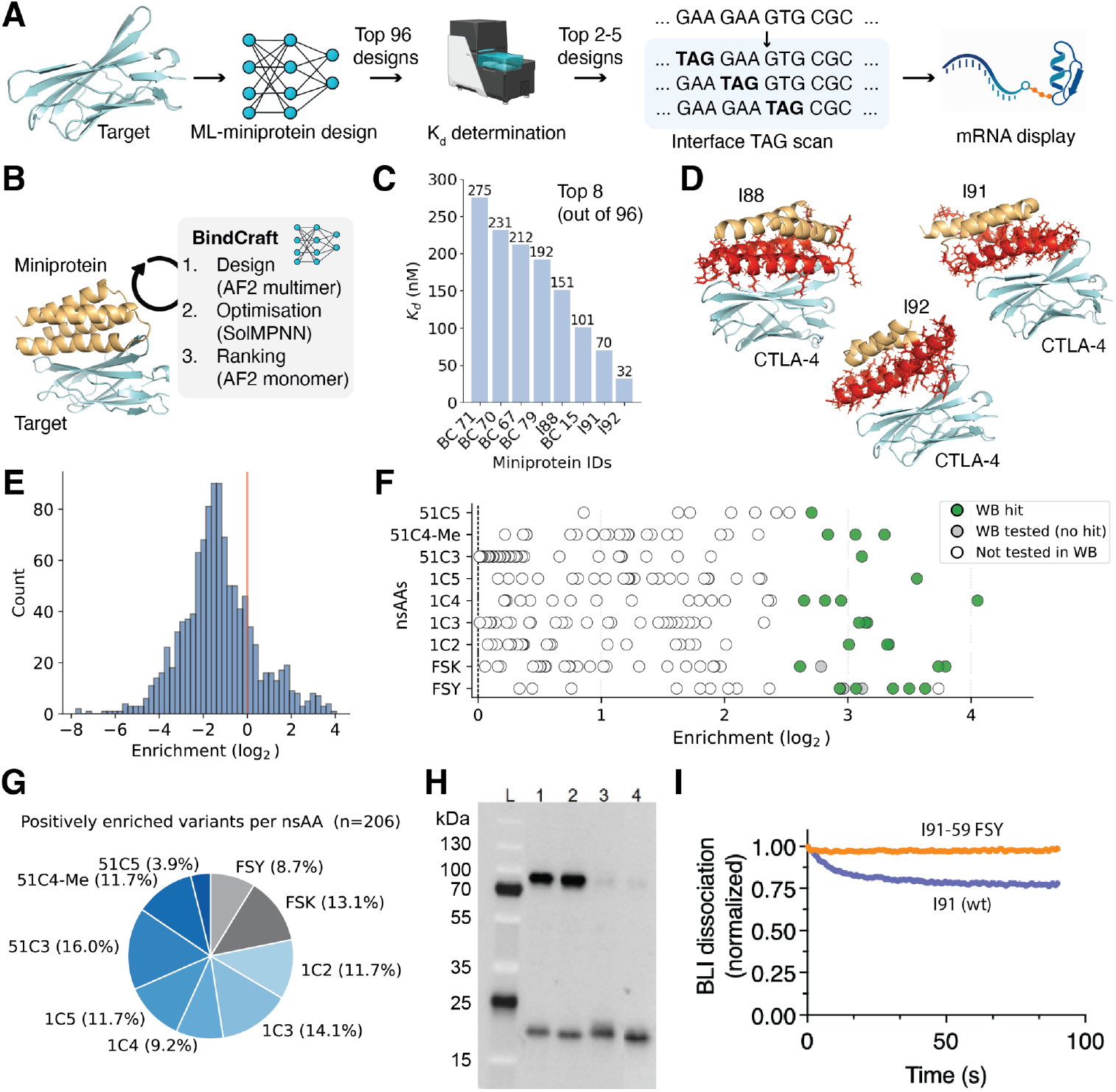
End-to-end discovery of covalent CTLA-4 miniproteins from generative design to durable engagement. **A**. Overall pipeline for discovering novel covalent binders: from target of interest to machine learning-driven *de novo* miniprotein design, top hit selection based on K_d_ measured by biolayer interferometry (BLI), and pooled mRNA display with covalent amino acid incorporation. **B**. BindCraft workflow: initial design with AlphaFold2 multimer, sequence optimization with soluble MPNN, and ranking with AlphaFold2 monomer confidence scores. **C**. Binding affinity values (K_d_, in nanomolar) of the top eight miniprotein designs (out of 96 tested). **D**. AlphaFold2 structure predictions of the three highest-affinity miniprotein designs. Orange and red: miniproteins. Red: miniprotein residues at the binding interface selected for UAG scanning. Cyan: target protein (CTLA-4). **E**. Enrichment distribution after selection. **F**. Positively enriched variants by nsAA. Green dots show positive hits in western blot (WB) and white dots non-hits in WB; white circles were not tested in WB. **G**. nsAA distribution in the positively enriched variants. **H**. WB showing covalent adduct formation between WNP 1C3-conjugated I92 covalent miniproteins and CTLA-4. 1: I92 P57[1C3], 2: I92 E59[1C3], 3: I92 A75[1C3], 4: I92 L77[1C3]. **I**. Dissociation rate as observed in BLI of miniprotein I91 with FSY at position 59 (orange) compared to wildtype I91 (violet) in pure express at the same concentration.

We applied BindCraft ^1^, a structure-guided generative modeling framework, to design miniproteins targeting a defined surface patch on CTLA-4 (Fig. 3B). To maximize the likelihood of functional engagement, we selected an epitope overlapping the clinically validated ipilimumab binding site (PDB ID: 5XJ3) ^39^. The resulting designs use different scaffolds and contact residues than 8GAB, providing an independent test of the platform on the same target. From this computational campaign, we advanced 96 top-ranked designs for experimental characterization. Each construct was fused to a TwinStrep tag and expressed in 1.5 µL PURExpress reactions. Binding affinities were quantified directly from crude reaction mixtures by BLI, eliminating purification bottlenecks (Supplementary Fig. 14). Overall, 35% of designs exhibited sub-micromolar affinity, with five miniproteins (BC79, I88, BC15, I91, and I92) achieving K_d_ values below 200 nM after a single round of computational design (Fig. 3C, Supplementary Fig. 14).

Next, we generated a UAG-scanning library for nsAA incorporation to convert three of the binders into covalent ones. We selected residues for the UAG scan sitting at the binding interface according to the predicted 3D structure (Fig. 3D), yielding 162 total UAG positions (54, 48, and 60 for I88, I91, and I92 respectively). Using our mRNA display platform, we incorporated nine fluorosulfate-containing nsAAs at these sites, generating 1,458 potential covalent variants (162 positions × nine nsAAs).

Scaffold-specific PCR primers were required for sequencing, precluding direct cross-scaffold normalization; we therefore report enrichment ratios (post/pre) within each scaffold, after restricting the analysis to variants with pre-selection frequency ≥ 0.0025 to avoid ratios inflated by rare reads (which also correlates with no expression in WB, Supplementary Fig. S19). Sequencing identified 34 position × nsAA combinations with log_2_ enrichment > 2.32 (>5-fold), spanning 19 unique positions across the three scaffolds (Fig. 3E). The strongest hits were I92 59[1C4] (log_2_ = 4.05), I92 59[FSK] (log_2_ = 3.80), and I88 23[FSY] (log_2_ = 3.73). The most reproducibly engaged site was I92 59, hit above 5-fold by five different nsAAs (FSK, WNP-1C2, WNP-1C3, WNP-1C4, WNP-51C4-Me); positions I92 57 and I91 58 were each engaged by four different chemistries. FSY accounted for 8.7% of all positively enriched variants and FSK 13.1%, with the remaining 78.2% distributed among the novel nsAAs—consistent with covalent engagement being driven primarily by geometric compatibility at these positions rather than by any single warhead (Fig. 3G).

To benchmark the sequencing readout against a biochemical standard, we cross-validated 42 variants from the filtered library by WB (Fig. 3F,H, Supplementary Figs. 15,16); 27 (64%) registered as covalent hits. Log_2_ enrichment correlated with WB signal (Spearman ρ = 0.72, p ≈ 1×10^-10^), and the relationship was especially clean at the top of the ranking: the 10 most-enriched WB-tested variants were all confirmed as covalent hits (100% precision), 18 of the top 20 were hits (90%), and across all 28 positively enriched WB-tested variants the cut achieved 86% precision and 89% recall (24/27 WB hits captured). These results provide an independent confirmation of the NGS–WB concordance reported for the 8GAB scaffold (Fig. 2B), with the relationship even tighter at the top of the ranking.

To quantify the impact of covalent engagement on binding durability, we measured the dissociation kinetics of I91 with FSY at position 59 by BLI. We normalized protein levels across samples by incorporating a fluorescent N-terminal label (Cy5-Met) via misaminoacylated initiator tRNA, which also let us directly visualize covalent adduct formation by in-gel fluorescence (Supplementary Fig. 20). Wild-type I91 miniprotein dissociated with measurable off-rates, whereas a warhead version, FSY at position 59, showed no detectable dissociation over the monitored timescale (Fig. 3I). Because covalent engagement decouples residence time from initial binding affinity, weakly binding designs that would be discarded on K_d_ alone could be rescued into durable engagers. This property relaxes the affinity threshold required at the design stage and expands the set of usable scaffolds beyond those accessible to non-covalent binders.

### A semisynthetic route from discovery to scalable manufacturing

Cell-free translation is well suited to discovery, but scaling it cost-effectively to manufacturing volumes remains challenging. To bridge the two, we developed a semisynthetic route that installs the warhead post-translationally on recombinantly expressed protein. A key advantage of de novo miniprotein design is the ability to generate lysine-free scaffolds. Because our nsAAs are built around a lysine core, the warhead module can be coupled directly onto a single reintroduced lysine via NHS-ester chemistry, assembling the full nsAA on the protein itself (Supplementary Fig. 17A).

After identifying optimal covalent positions through mRNA-display screening, we reintroduced a single lysine at the top hit positions (e.g., 8GAB K51) to create precursor variants amenable to scalable semisynthetic production ^23^. These constructs were expressed and purified using standard recombinant workflows, after which covalent warheads were installed via activated-ester chemistry (e.g., WSP 51C4-Me NHS-ester) with a TEV-cleavable N-terminal tag absorbing α-amine labeling that was removed after coupling. The resulting semisynthetic covalent proteins gave clean HPLC profiles and single band by SDS-PAGE and retained proximity-dependent reactivity comparable to proteins produced directly by cell-free translation with chemically charged tRNA (Supplementary Fig. 17)

We demonstrated this strategy could also be extended to scaffolds containing native lysines, including nanobodies and scFvs, by globally substituting lysines to arginines. This preserved binding affinity across scaffold types, with K_d_ values remaining in the nanomolar range (Supplementary Fig. 21A). One exception was observed in Porustobart (heavy-chain antibody), where structural analysis suggests K32 may interact with CTLA-4 E59 at the binding interface (Supplementary Fig. 21B). Reintroducing a single lysine at a covalent hotspot would then allow site-specific warhead installation on scaffolds otherwise inaccessible to homogeneous chemical modification. Together with retrosynthetically designed lysine-based nsAAs, this provides a route from mRNA display hits to scalable semisynthetic production across diverse scaffold types.

### Beyond covalency: dual nsAA incorporation and fluorogenic warheads

As AminoX decouples tRNA charging from synthetase constraints, any nsAA can be paired with any orthogonal tRNA, and multiple charged tRNAs can be used in the same reaction to decode distinct codons. This opens two strategies for multifunctional proteins: incorporating multiple different nsAAs at separate positions, or incorporating a single nsAA that combines multiple functions (Fig. 4C).

**Fig. 4.**
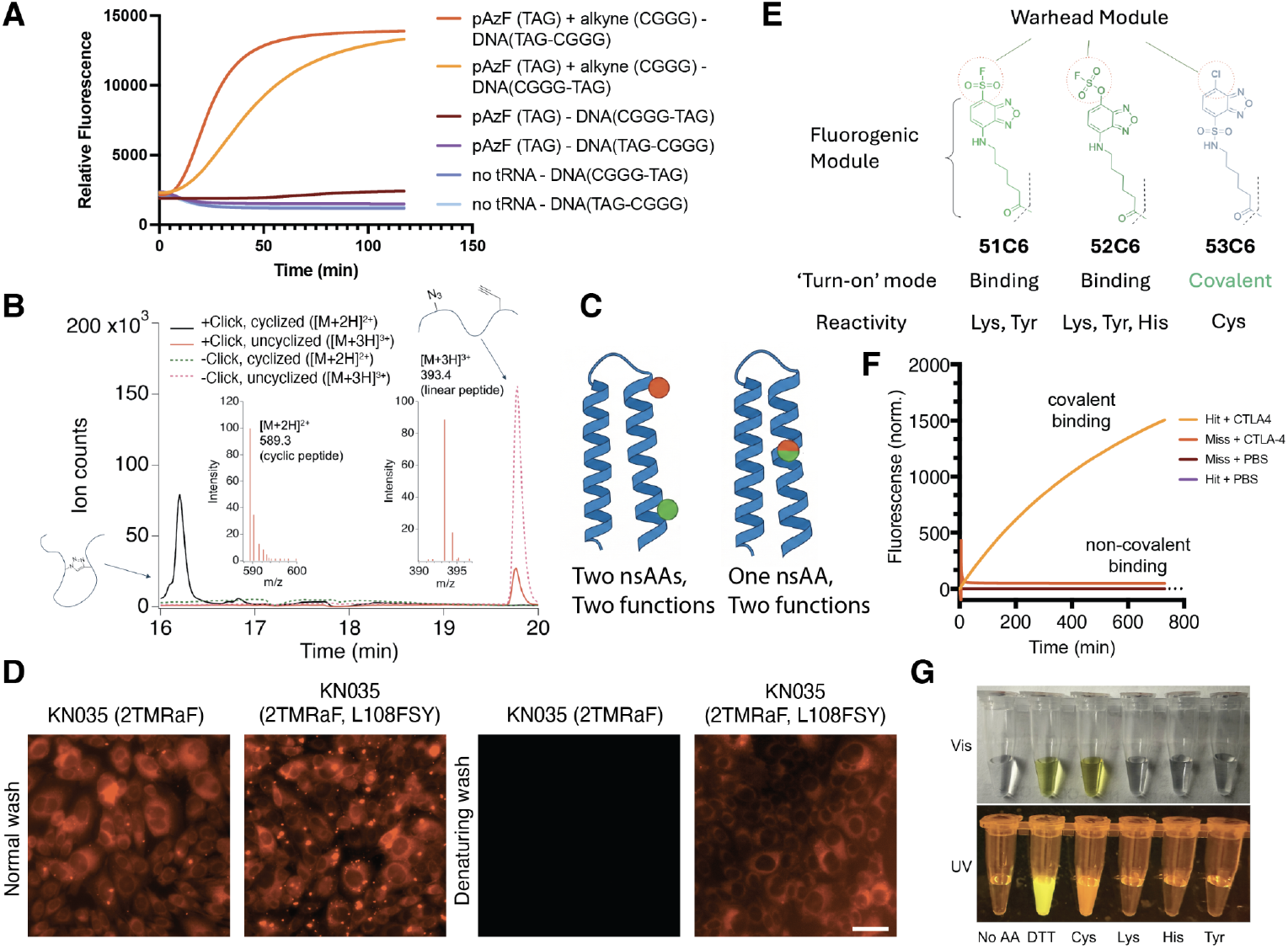
Multifunctional protein engineering through dual nsAA incorporation and fluorogenic covalent warheads. **A**. GFP reporter assay for dual nsAA incorporation. Engineered sfGFP templates contain TAG (amber) and CGGG (quadruplet) codons at positions 2 and 5 DNA(TAG-CGGG) or positions 3 and 8 DNA(CGGG-TAG). **B**. HPLC-MS of pAzF/pPaF-containing peptide before (dashed) and after (solid) Cu(I)-catalyzed click chemistry. Traces show retention time and observed charge states ([M+3H]^3+^ linear; [M+2H]^2+^ macrocycle). **C**. Schematic of two strategies for multifunctional protein engineering. Left: Dual nsAA incorporation, where two distinct nsAAs (covalent and fluorescent/click-chemistry) are site-specifically installed at different positions using orthogonal codons. Right: Single multifunctional nsAA with integrated dual functionality (covalent warhead and fluorogenic reporter) at one position. **D**. Dual nsAA incorporation enables fluorescent tracking of covalent target engagement on fixed CHO cells (scale bar = 40 µm). **E**. Fluorogenic covalent nsAA designs combining NBD chromophore with distinct warhead modules. 51C6 (sulfonyl fluoride) and 52C6 (fluorosulfonyloxybenzoyl) exhibit binding-activated turn-on. 53C6 integrates the warhead and fluorogenic modules into a single NBD-Cl group where covalent bond formation with thiols activates fluorescence. **F**. Fluorescence analysis of 8GAB miniprotein incorporating WSP-51C4-Me. Hit position 8GAB I59[51C4-Me] shows time-dependent fluorescence increase upon incubation with CTLA-4; miss position 8GAB Y22[51C4-Me] and PBS controls show no increase. **G**. Covalent-activated fluorescence of 53C6 is colorless and non-fluorescent. Reaction with thiols (cysteine or DTT) turns it yellow and strongly fluorescent under UV.

To demonstrate dual nsAA incorporation, we engineered an sfGFP reporter with both a TAG and a quadruplet (CGGG) codon in positions required for fluorescence, such that full-length GFP is produced only when both codons are successfully decoded. Co-incorporation of p-azido-L-phenylalanine (pAzF, at TAG) and p-propargyloxy-L-phenylalanine (pPaF, at CGGG) produced robust fluorescence, with negligible background in single-tRNA or no-tRNA controls (Fig. 4A).

We tested two applications that dual incorporation enables: post-translational macrocyclization, and fluorescent tracking of covalent target engagement on cells. First, we generated macrocyclic peptides through Cu(I)-catalyzed click chemistry between incorporated pAzF and pPaF residues. HPLC-MS confirmed cyclization through retention-time and charge-state shifts consistent with the more compact macrocyclic structure (Fig. 4B), establishing that both nsAAs were chemically intact after incorporation and accessible to post-translational modification. Second, we used dual incorporation to track covalent target engagement on cells. Traditional immunofluorescence workflows for covalent binders require purification followed by chemical conjugation with a fluorescent probe ^33^; in contrast, cell-free translation enabled direct production of a dual-modified PD-L1 nanobody ^40^, KN035 (2TMRaF, L108[FSY]), integrating a red fluorescent amino acid (tetramethylrhodamine-aminophenylalanine, TMRaF) at position 2 via a CGGG codon and the covalent warhead FSY at position 108 via TAG. Fixed CHO cells expressing PD-L1 were stained with KN035 (2TMRaF) (non-covalent control) or KN035 (2TMRaF, L108[FSY]) (covalent). Both nanobodies labelled cells after gentle wash (left). After a denaturing wash, only the covalent variant remained bound (right), confirming stable covalent target engagement (Fig. 4D).

For multifunctional nsAA design, we drew on the solvatochromic nanosensor framework established in our prior work ^23^, to first design a series of fluorogenic covalent NBD-SF amino acids (Fig. 1F, WSP-51C3, WSP-51C4-Me, WSP-51C5). These combine a sulfonyl fluoride warhead with the NBD chromophore, whose fluorescence increases upon environmental changes such as burial in a binding interface. We reasoned that covalent engagement would position the NBD chromophore in a fluorescence-enhancing environment. Incorporation of WSP-51C4-Me at 8GAB I59, a covalent hit from our CTLA-4 mRNA display screen, produced a time-dependent fluorescence increase upon CTLA-4 addition, while a non-hit position 8GAB Y22 and PBS controls showed no signal (Fig. 4F). Fluorogenic variants retained full CTLA-4 blocking activity in cell-based assays (Fig. 2E), indicating that the optical readout does not compromise therapeutic function.

To extend residue-level reactivity, we designed two additional fluorogenic warheads (Fig. 4E). WSP-52 series replaces the directly-attached sulfonyl fluoride of the 51 series with an aryl-spaced fluorosulfonyloxybenzoyl group, retaining binding-activated turn-on while extending reactivity to histidine. The WSP-53 series collapses warhead and fluorogenic modules into one NBD-Cl group where thiols, e.g., cysteine, displaces the chloride to form the covalent bond and turn on fluorescence in the same step (Fig. 4G). Together, these three new fluorogenic warhead nsAA classes give complementary residue coverage (Lys, Tyr, His, Cys). Ribosomal incorporation was confirmed by GFP reporter assay for 52C6 (Supplementary Fig. 22) and by HPLC-MS for the WSP-53 series (Supplementary Fig. 23). Cys reactivity of 53C6 was confirmed by HPLC-MS (Supplementary Fig. 24). These fluorogenic warheads provide a fluorescence readout that could replace WB or MS for primary validation in larger screens.

Together, these results show that AminoX extends from covalent engagement to a broad toolkit for multifunctional protein engineering. Dual incorporation enables spatial combination of orthogonal chemistries within a single protein; integrated multimodal nsAAs collapse those modalities onto a single residue.

## Discussion

AminoX establishes a general platform for incorporating chemically diverse non-standard amino acids (nsAAs) into folded proteins at scale. By eliminating the need for evolved aminoacyl-tRNA synthetases, the platform uncouples nsAA chemistry from protein-engineering timelines, turning warhead design, linker geometry, and incorporation site into three independent axes of optimization rather than a sequence of engineering problems.

This uncoupling enables a mode of discovery that has been out of reach for covalent protein engineering. Rather than characterizing warhead–position combinations one at a time, AminoX screens hundreds in a single pooled selection, with sequencing enrichment reliably pre-filtering covalent hits for targeted biochemical validation. In the CTLA-4 screens alone, this compressed what would have been more than 2,000 individual synthesis-and-assay cycles.

Using BindCraft-designed miniproteins generated from only the target structure as input, we identified covalent variants with enhanced functional activity in *in vitro* assays. Because our warheads are retrosynthetically designed from lysine, hits from pooled selection can be reformatted as activated-esters for site-specific installation on recombinantly produced protein, providing a direct path from discovery to scalable manufacturing.

A further consequence of decoupling residence time from initial binding affinity is that covalent engagement reshapes the design constraints for selectivity. Selectivity between closely related target variants—for example, a disease-associated mutant and a wild-type homolog expressed in healthy tissue—is conventionally limited by the affinity differential between the two, which can be small when the variants differ by only a few residues. Covalent engagement could provide an additional discriminator: a warhead positioned to engage a residue present in one variant but not the other can convert a modest non-covalent preference into a durable, irreversible interaction with one variant while the other dissociates normally. This could prove useful for targeting disease-associated variants of otherwise poorly differentiated proteins, including oncogenic point mutants and pathogen escape variants.

Beyond covalency, the same direct-acylation foundation supports arbitrary nsAA combinations, enabling multifunctional proteins that carry covalent warheads, fluorogenic reporters, or click handles—either at separate positions or integrated into a single residue. Dual incorporation and integrated multimodal nsAAs both derive from the same underlying capability: efficient direct tRNA acylation makes arbitrary nsAAs accessible without evolved synthetases.

Several limitations of the current work bear mention. First, the workflow demonstrated here applies to biologics produced ex vivo, either by cell-free translation or by recombinant expression followed by semisynthetic warhead installation. Cell-surface display of covalent binders, such as covalent chimeric antigen receptors (CARs) for adoptive cell therapy, would require extending direct-acylation chemistry to living cells. Second, while we validated mRNA display compatibility across nanobodies, scFvs, and heavy-chain antibodies, covalent selection itself was only run on miniprotein scaffolds; extending pooled selection to larger scaffolds remains to be demonstrated. Third, our warhead panel is dominated by SuFEx-based chemistries; broader electrophile classes like Michael acceptors, α-haloacetamides, and activated esters are compatible with direct acylation in principle, but untested here. Fourth, scaling beyond the warheads tested here will require expanded barcode sets, deeper sequencing, and validation that pooled cross-talk remains negligible.

AminoX treats the genetic code as a programmable interface for chemistry rather than a fixed constraint, bringing folded-protein engineering closer to the design freedom that medicinal chemists have long enjoyed with small molecules. As generative design continues to expand the set of accessible binder scaffolds, coupling those scaffolds to arbitrary chemistry offers a route to biologics that can be engineered, not merely evolved.

## Acknowledgements

We thank Samuel Inverso, Steve Liapis, Elias Quijano, Jenny Tam, and Jim Niemi for helpful discussions about this project. We thank Ken Carlson and Sylvie Bernier for advice in target selection and cell assays.

## Author contributions

H.P., E.K., M.M. designed and performed experiments, analyzed the data, and wrote the manuscript. A.F., S.K., C.E., D.S., S.R. performed experiments and edited the manuscript. J.J.C. and G.M.C. directed overall research and edited the manuscript.

## Data and materials availability

All data needed to evaluate the conclusions in the paper can be found in the paper and/or the Supplementary Materials. Sequences used in this study are listed in Table S1 of the Supplementary Materials. Correspondence and requests for materials should be addressed to G.M.C. and J.J.C.

