## Supplementary Materials for "Genetic code expansion enables programmable covalent protein design"

**This PDF file includes:**

Materials and Methods
Supplementary Figures (S1 to S24)
Supplementary Tables (S1)
Supplementary References

### Materials and Methods

**Reagents:** Stock solutions were prepared in anhydrous DMSO avoiding prolonged exposure to the room temperature and stored in -80ºC. 5'-Phospho-2'-deoxyribocytidylylriboadenosine (pdCpA) was purchased from Dharmacon. Esterification of Boc-nsAA-OH to pdCpA, deprotection of Boc-nsAA-pdCpA and ligation to tRNA(-CA) were performed following protocols [1]. 1-Step Ultra TMB-ELISA Substrate Solution (Cat. No. 34028) was purchased from Life Technologies. DNA was purchased from IDT.

**Antigens**: The receptor binding domain (RBD) antigens were purchased from Genscript: SARS-CoV-2 Spike RBD, CoV19 (Cat. No. Z03491) and PD-L1-Fc was purchased from Peprotech (Cat. No. 310-35-500UG). CTLA-4-Fc (Cat. No. CT4-H5255-200ug), Biotinylated CTLA-4 ( Cat. No. CT4H82F325UG), mouse Recombinant CTLA-4 Fc Tag (Cat No. 103014-302), human CD28-Fc (Cat. No. CD8-H525a-100ug) and mouse CD-28-Fc (CD8-M5258-100ug) were purchased from Acro.

**Antibodies:** Ipilimumab (Cat. No. LT1600-1MG), Goat anti-Human IgG (H+L) Secondary Antibody, HRP (Cat. No. A18805), Goat anti-Human IgG Fc Cross-Adsorbed Secondary Antibody, DyLight 650 (Cat. No. SA510137) and DYKDDDDK Tag Monoclonal Antibody (FG4R), HRP (Cat. No. MA191878HRP) were purchased from Life Technologies. Monoclonal anti-FLAG M2-FITC (Cat. No. F4049-.2MG) was purchased from Sigma Aldrich. Control antibodies Anti-CTLA-4 (JA1020) and Anti-PD-L1(Cat.# J1201) were purchased from Promega Corporation. Anti-ALFA-Tag antibody (HRP, Cat. No. ABIN7272969), and anti-ALFA-Tag antibody (Alexa Fluor 647, Cat. No. ABIN7272960) were purchased from Antibodies-Online Inc. Strep Tactin DY-649 (Cat. No. 502100145) was purchased from Fisher. Strep Tactin HRP conjugate (MSPP-21502001, EA) was purchased from VWR.

**Materials:** BLI Strep-Tactin XT Probes (Cat. No. 160033), Streptavidin (SA) probes (Cat. No. 160002), plates (130161-10Pcs) and buffers (Cat. No. 120063) were purchased from Gator Bio.

**Cell lines and cell assays:** CHO-K1 cells expressing PD-L1 (Cat. No. M00543) and CHO cells expressing CTLA-4 (Cat. No. M00530) were purchased from Genscript, cultured in Ham's F-12K (Kaighn's) (Cat. No. 21127-022, Life Technologies), 10% FBS (Cat. No. 10099-141, Life Technologies).and tested routinely for mycoplasma contamination. Thaw and use MOA CTLA-4 Promega bioassays (Cat. No. JA3001) were purchased from Promega and used following the manufacturer's instructions with an incubation time of 8h.

**Miniprotein Design:** Miniprotein binders were computationally designed using BindCraft (https://github.com/martinpacesa/BindCraft), a structure-based protein design platform. The

design protocol involved sampling over a period of time. Design candidates were filtered using BindCraft's built-in quality metrics as acceptance criteria. This initial filtering yielded 134 designs that passed the BindCraft quality threshold.
We selected 96 top candidates from the 134 passing designs based on their average ipTM (interface predicted Template Modeling) scores. The average ipTM metric evaluates the predicted accuracy of the protein-protein interface, with higher values indicating greater confidence in the designed binding interface. The 96 highest-scoring designs by this metric were designed into DNA eBlocks by including a 5' T7 followed by a RBS, and 3' FLAG peptide and T7 terminator; and advanced for binding validation with bio-Layer Interferometry (BLI).

**Software**: Graphs were plotted and analyzed in Prism 10, GraphPad Software, www.graphpad.com. Biolayer interferometry data was analyzed in the Gator analysis software.

**Synthesis of proteins in cell-free protein synthesis (CFPS) systems:** DNA templates for the in vitro transcription and translation reactions contained optimized sequences of a T7 promoter, a Shine-Dalgarno sequence, the open reading frame with an in-frame amber stop codon (* = TAG) that would be suppressed in the presence of charged full-length tRNA species. CFPS reactions were carried out using PURExpress® Δ RF1 or NEBExpress Cell-free E. coli Protein Synthesis System (NEB, Ipswich MA) following the manufacturer's instructions and supplying 20ng/µL DNA templates with nsAA-tRNA$_{CUA}$ (at final ~8 µM), and 1.5 units/µL RNase Inhibitor Murine (NEB). Reactions (1.5–50 µL) were incubated in 0.2 mL thin wall PCR tubes (Thermo Fisher) at 30–37 °C for 60min–300min. 1.5 µl of CFPS extracts containing proximity-dependent covalent proteins were directly mixed with 0.75 µl of 0.5-1mg/ml purified protein targets and incubated for 6h at 37ºC to probe for covalent binding.

**Gel electrophoresis:** Reactions were analyzed by running 2 µL of the reaction with 2X loading buffer (Novex) and 10X reducing buffer (Novex), followed by heating to 90ºC for 10min to denature proteins and dissociate protein complexes that were not covalently attached. We ran the samples in parallel with 10 µL Precision Plus Protein™ fluorescent protein ladder (Bio-Rad) or PAGE Ruler Plus (PI26619, Invitrogen) in 1.0mm Invitrogen™ Novex™ WedgeWell™ 10–20%, Tris-Glycine mini protein gels (Thermo Fischer) following the manufacturer's instructions and materials. Electrophoresis was performed in 1× Novex running buffer (Tris-Glycine-SDS) at 120–150 V for 50min.The in-gel fluorescence was measured using a Biorad Gel Doc XR+ Imaging System and the gels were Coomassie stained with InstantBlue protein stain (Novus Biologicals), following the manufacturer's instructions. The images were quantified by ImageJ.

**Western blot:** PAGE gels were washed, imaged and transferred to PVDF iBlot membranes and transferred in an iBlot machine for 6min, according to the manufacturer's protocol. Membranes were washed and blocked for 1 h at room temperature in a blocking buffer (5% non-fat dry milk

-Rockland- in PBS-T; PBS pH 7.4, 150 mM NaCl, 0.05% Tween-20). Blots were washed and incubated with two simultaneous secondary antibodies diluted in the blocking buffer (1:5000 for fluorescent anti-human Fc antibodies, and–1:50000 for anti-FLAG-HRP; anti-StrepTactin-HRP or anti-ALFA-HRP) for 1h at room temperature, followed by three additional PBS-T washes. Signals were developed using enhanced chemiluminescence (ECL) substrate (ClarityMax, BioRad) and imaged on a Biorad Gel Doc XR+ Imaging System. Band intensities were quantified using ImageJ.

**Biolayer Interferometry:** Assays were performed on a Gator® BLI instrument at 25 °C following the manufacturer's guidelines. All reagents were equilibrated to room temperature, and samples were dispensed into a black 96-well plate, or 384 tilted-bottom black plate (typically 60-200 µL per well). Biosensors were pre-equilibrated in an assay buffer for 10 min to hydrate and establish a baseline, we regenerated all StrepTactinXT sensors up to 15 times, following manufacturer's advice. In a typical experiment, sensors were dipped into wells containing CFPS-expressed proteins (20-40X diluted from PURExpress in the assay buffer, PBST) for 5–10 min. For kinetic measurements, sensors were first equilibrated in assay buffer (baseline, 60s), then transferred to wells containing analyte at different concentrations (typically between 0-2.5µM, association phase, 180–600 s), followed by transfer back into buffer-only wells (dissociation phase, 300–1200 s). Parallel reference sensors (e.g., unloaded or loaded with negative control CFPS no protein) were included to control for non-specific binding and drift. Raw sensograms were reference-subtracted and aligned using the Gator® analysis software. Association and dissociation curves were globally fit to a 1:1 binding model (or alternative models as appropriate) to extract association (ka), dissociation (kd) rate constants and equilibrium dissociation constants (KD = kd/ka). Only datasets with high signal-to-noise ratio and good fit statistics (e.g., $\chi^2$ within software-recommended thresholds) were included in final analyses.

**mRNA/cDNA display:**: ORF libraries for mRNA/cDNA display were constructed stepwise. The CTLA-4 8GAB and I88/I91/I92 libraries containing in-frame TAG stop codons were pooled from individual eBlocks ordered from IDT (Supplementary Table S1). Our mRNA/cDNA display approach involves translating nsAA-containing protein binder libraries and covalently linking them to their cDNA via a puromycin linker using our previously optimized protocol [1] with the following modifications for covalent warhead workflow: CFPS reactions using PURExpress® ΔRF1 contained 8 µM nsAA-tRNACUA and were performed at 30°C for 90 min. CTLA-4 was added to 50 µg/mL final concentration and incubated at 37°C for 90 min or durations indicated in main text, followed by 7:1 addition of quenching solution (40 mM L-tyrosine, 200 mM L-histidine in 100 mM Tris pH 8.5) and incubation at 50°C for 15 min then 37°C for 15 min. Reverse transcription was performed at 37°C for 1 hr using ProtoScript® II Reverse Transcriptase (NEB). For barcoded competition experiments, each nsAA library was reverse transcribed with a unique 8-bp barcoded primer (Supplementary Table S1) before pooling. Stringent selection included incubation at 70°C for 15 min to denature non-covalent complexes, followed by streptavidin pulldown and washes with 5M guanidine. Post-selection DNA was amplified using construct-specific forward primers (P3-P6) paired with universal reverse primer P7, then sequenced by NGS (Amplicon-EZ, Azenta/Genewiz).

**Illumina next-generation sequencing (NGS) data analysis and read counting:** NGS-based amplicon sequencing was performed using the Amplicon-EZ service of Azenta/Genewiz and DNA amplicons from each selection cycle. cDNA in samples from washes and elutions were amplified by Pri5-Pri6 and were prepared following Amplicon- EZ sample submission guidelines. Raw reads (236 bp paired-end) were merged using BBMerge, and filtered for Phred quality scores at or above 20. Resulting reads were forwarded and trimmed using a custom Python script, which identified the first 18 bp of the constant region. The read frequency was calculated as the fraction of each unique sequence divided by the total
number of trimmed sequences detected within a sample. Approximately 200 unique reads were identified in all samples from the mRNA/ cDNA display evolution rounds, and fold enrichment was calculated by dividing the read frequencies of subsequent rounds by the read frequency identified in the original library.

**Immunofluorescence microscopy and image acquisition:** For microscopy experiments leading to Fig. 5D, KN035_2TMRaPhe, L108FSY, and KN035_2TMRaPhe were expressed in PURExpress. CHO-K1 and CHO-K1 cells expressing PD-L1 were cultured in 96-well plates overnight, fixed in 4%PFA, washed in PBS and cells were stored in the same buffer at 4C for up to 1 week before staining and imaging. At the day of imaging, the media was exchanged into 1 x PBS + protein extracts, and directly imaged. Phase and fluorescence images were acquired using a Nikon Ti2 Eclipse inverted microscope equipped with a Plan Apo Lambda 20X (0.75 NA, DIC N2) oil objective and Andor Zyla sCMOS camera. NIS-Element AR software was used for image acquisition. Image processing was performed in FIJI. Images were scaled, cropped and rotated without interpolation. Linear adjustment was performed to optimize contrast and brightness of the images. Figure construction was performed in Adobe Illustrator.

**Mass spectrometry**. For peptide HPLC/MS, equal volume of 2% formic acid was added to PURE reactions to precipitate large proteins. Samples were then centrifuged at > 12,000 x g for 10 min. Samples in 1% formic acid (1 µL injection) were ran on an Agilent 1290 UPLC using a Poroshell 120 SB-Aq column (2.7 µm, 2.1 × 50 mm; Agilent) with a linear gradient from 5% to 100% acetonitrile over 3.5 min at a flow rate of 0.6 mL/min with 0.1% formic acid in the mobile phase. Mass spectra were acquired using an Agilent 6530c QTOF with the following source and acquisition parameters: Gas temperature = 300 ºC; gas flow = 8 l/min; nebulizer = 35 psig; capillary voltage = 3500 V; fragmentor 175 V; skimmer 65 V; oct 1 RF vpp = 750 V; acquisition rate = 3 spectra/s; acquisition time = 333.3 ms/spectrum; collision energy 0 V. Extracted ions for NT-formyl peptides (fM[NBDxK]PVFV and fMFPV[NBDxK]V; [M+H]+ m/z = 1024.4920) were monitored within a 10 ppm window.

**tRNA ligation and quantification.** The enzymatic esterification of tRNA(-CA) species to nsAA-pdCpAs (resulting in nsAA-tRNACUA) was done as previously described [1]. Briefly, 500 µg of PylT tRNA(-CA)$_{CUA}$ or Mycoplasma capricolum Trp1 tRNA(-CA)$_{CUA}$ was dissolved in 625 µL 10 mM HEPES + 2.5 mM $MgCl_2$ and folded by heating to 95 °C for 3 min with a subsequent gradual cool-down to 25°C over 20 min. The aminoacylation reaction to obtain the full length nsAA-tRNA$_{CUA}$ contained the final concentrations of 300 µg/mL folded tRNA(-CA), 0.3 mM nsAA-pdCpA (from 3 mM DMSO stock), 1 X of T4 RNA Ligase buffer (from 10x, NEB), 0.125 mM ATP and 600 units/mL of T4 RNA Ligase 1 (NEB). This reaction was incubated at 4°C for 2 h. The nsAA-tRNA CUA was extracted with acidic phenol chloroform (5:1, pH 4.5), ethanol precipitated, washed, air-dried and stored at −80°C.

**N-terminal fluorescent labeling for expression normalization.** For quantitative comparison of protein variants, *E. coli* initiator tRN (fMet) was misaminoacylated with a Cy5-labeled methionine analog using an approach adapted from [2]. Methionine-pdCpA was conjugated to Cy5-NHS ester at the α-amino group and ligated to truncated initiator tRNA using T4 RNA ligase. The resulting Cy5-Met-tRNA(fMet) was added to PURExpress reactions alongside nsAA-charged suppressor tRNAs. Because Cy5-Met incorporation reflects nascent protein synthesis, expression levels were quantified directly by in-gel fluorescence imaging of SDS-PAGE gels.

For competitive dissociation experiments (Fig. 3I), covalent and non-covalent miniprotein variants were expressed in separate PURExpress reactions containing 8 µM Cy5-Met-tRNA(fMet). Different volumes of each reaction were loaded on SDS-PAGE to determine the amount needed for equal protein concentration based on Cy5 fluorescence intensity. Proteins were then mixed with CTLA-4 (0.5 mg/mL, 1 µL) at defined ratios and incubated for 12 hours at 37°C to allow covalent bond formation. Samples were analyzed by BLI for dissociation kinetics and by gel electrophoresis with in-gel fluorescence detection for covalent adduct formation.

### Supplementary Figures

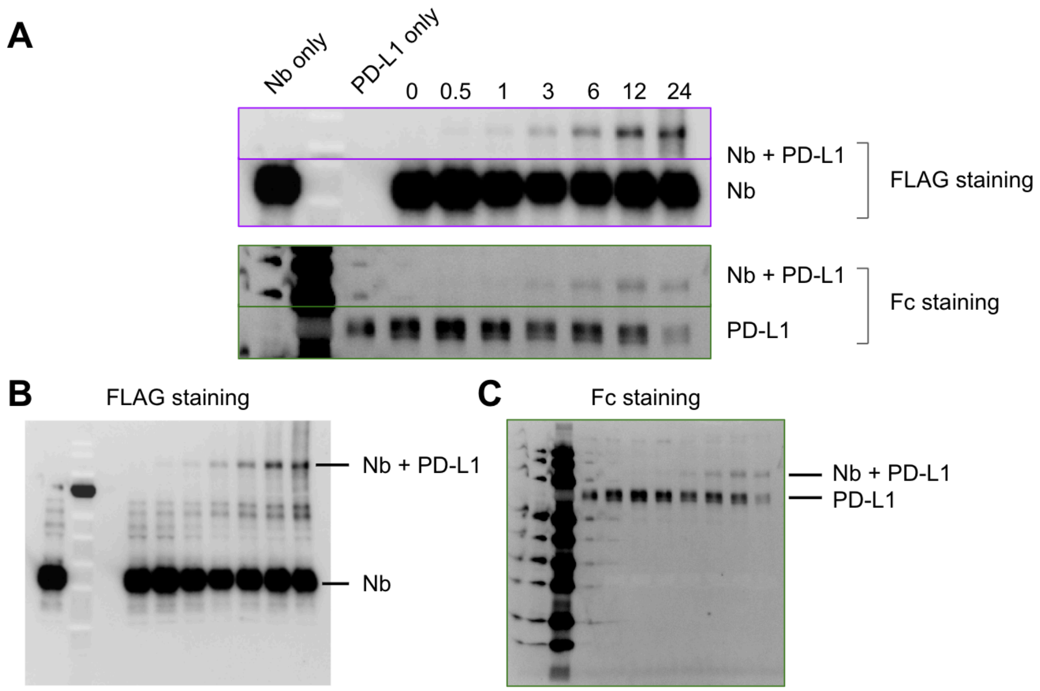


**Fig. S1 | Western blot with double staining is used to verify co-localization of FSY-containing covalent proteins bound to their targets in covalent-binding time-courses.** 1.5 µl of FSY containing proteins were expressed by CFPS for 4h at 30C followed by incubation for increasing time with 0.75 µl of PD-L1-Fc. Gels were stained with fluorescent anti-human IgG and anti-FLAG-HRP. **A.** Results summary. **B.** Full gel with anti-FLAG staining. **C.** Full gel with anti-human IgG staining.

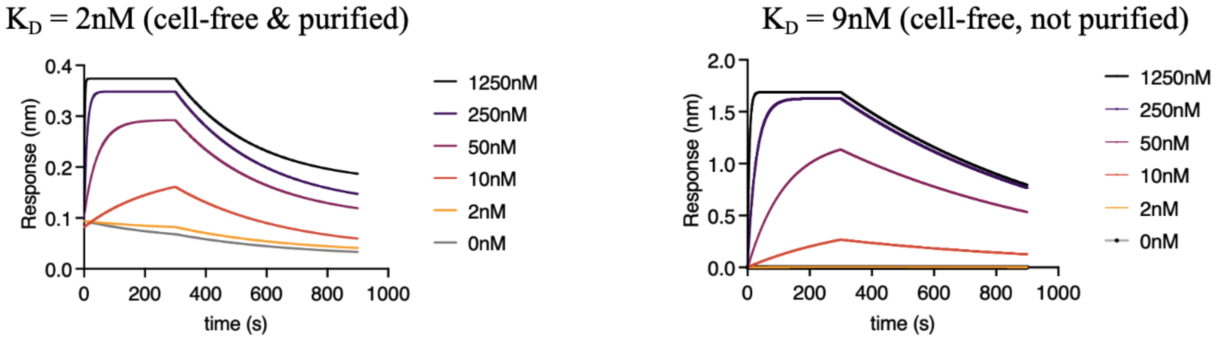


**Fig. S2 | Biolayer interferometry data of PD-L1 nanobody binding before and after purification.** KN035 expressed in cell-free with a His-tag was loaded onto Ni-NTA tips and assayed for PD-L1 binding before and after purification. Measured $K_d$ is comparable to previously published values [3].

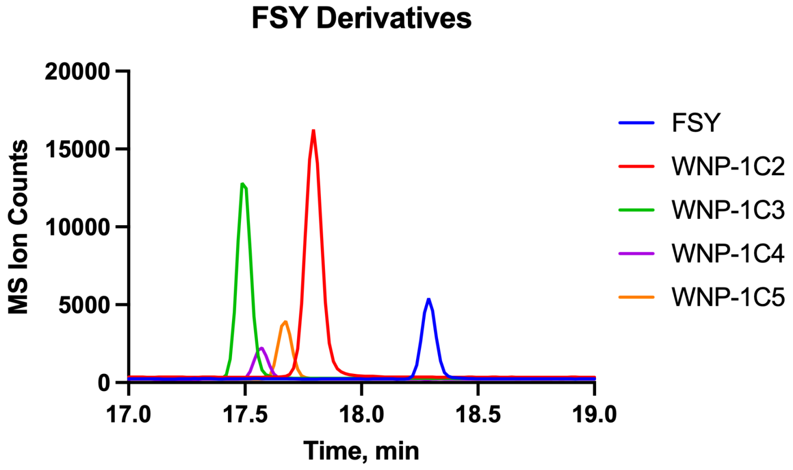


**Fig. S3 | Mass spectrometry verification of site-specific ribosomal incorporation of fluorosulfate-containing amino acids.** FSY and chain-extended variants (WNP-1C2, WNP-1C3, WNP-1C4, and WNP-1C5) were incorporated into position * of the peptide fMFPV*V. Raw ion counts shown as a function of time.

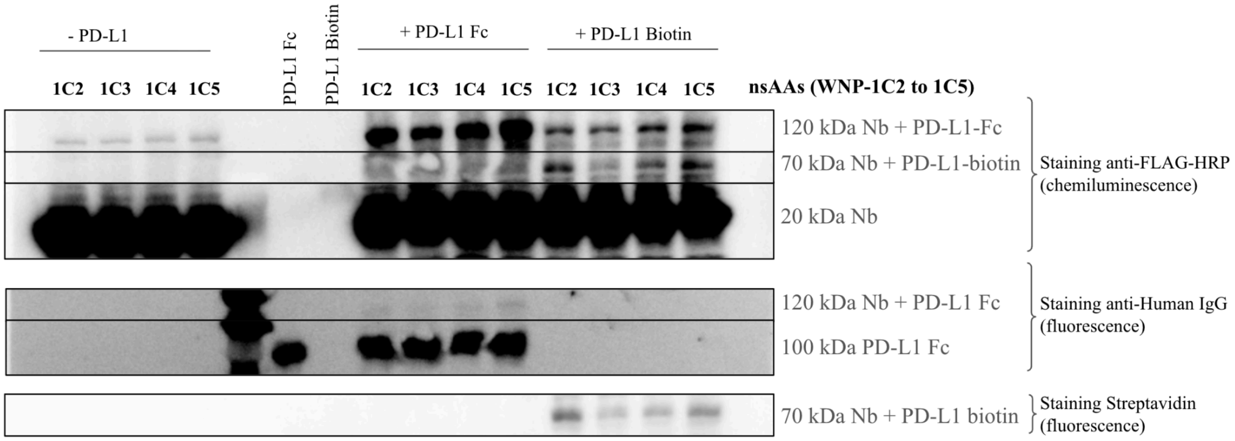


**Fig. S4 | Western blots to identify new covalent proteins containing amino acids of different lengths that bind PD-L1.** 1.5 µl of proteins containing the nsAA (WNP-1C1, WNP-1C2, WNP-1C4 and WNP-1C5) at position T110[X] were expressed by CFPS for 4h at 30C followed by incubation for 6h 0.75 µl of PD-L1-Fc, PD-L1-Biotin, or negative control. Gels were stained with fluorescent anti-human IgG-Fluor (stains PD-L1 Fc), anti-FLAG-HRP (stains cell-free expressed KN035 Nanobody) and Streptavidin-Fluor (stains biotinylated PD-L1). Nb stands for nanobody KN035. After covalent interaction, new covalent adducts are observed at the positions that correspond to added molecular weights of the nanobody plus PD-L1.

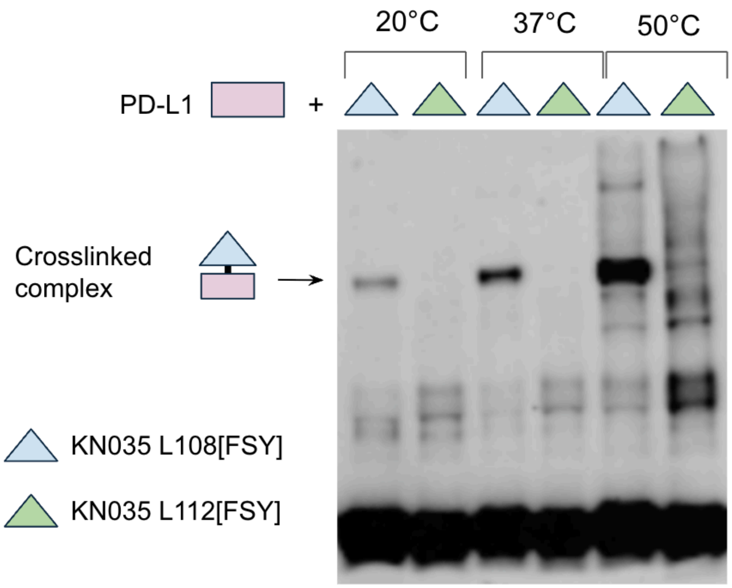


**Fig. S5 | Effect of temperature on covalent adduct formation and specificity.** Covalent nanobodies (KN035 L108[FSY] & KN035 S112[FSY]) incubated with PD-L1 for four hours at different temperatures. Higher temperatures increase warhead reactivity but decrease specificity, with non-specific crosslinking to cell-free protein synthesis components observed at elevated temperatures.

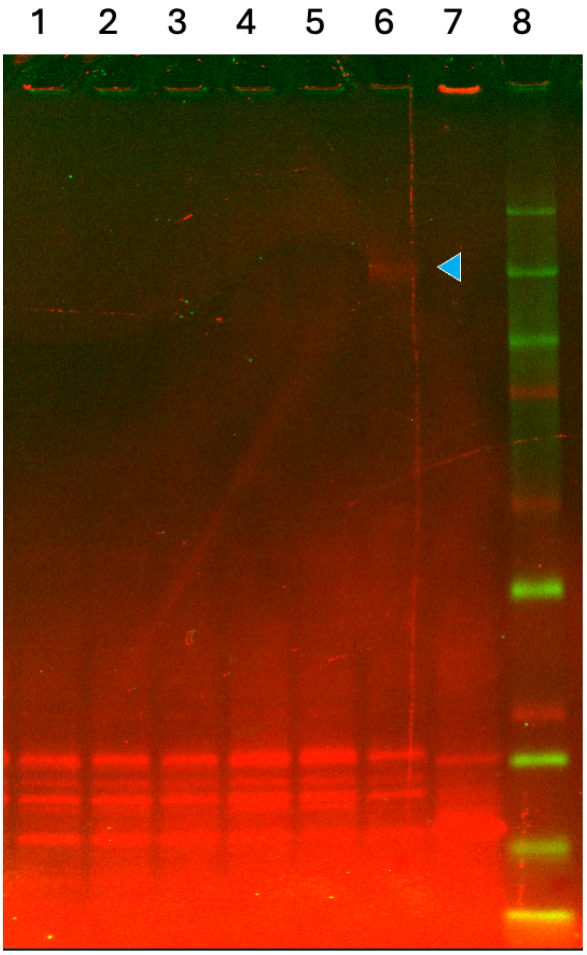


**Fig. S6 | Single-round mRNA display selection enriches warhead-specific covalent binders from a pooled TAG scan library.** Pooled TAG scan library of 8GAB was translated with FSY, selected against CTLA-4, and enriched DNA re-translated with indicated covalent nsAAs. Lanes 1–3: PBS control; Lanes 4–6: CTLA-4; Lane 7: no nsAA; Lane 8: ladder. The arrow indicates covalent adduct observed only with FSY + CTLA-4 (lane 6), confirming warhead-specific enrichment after one round of selection.

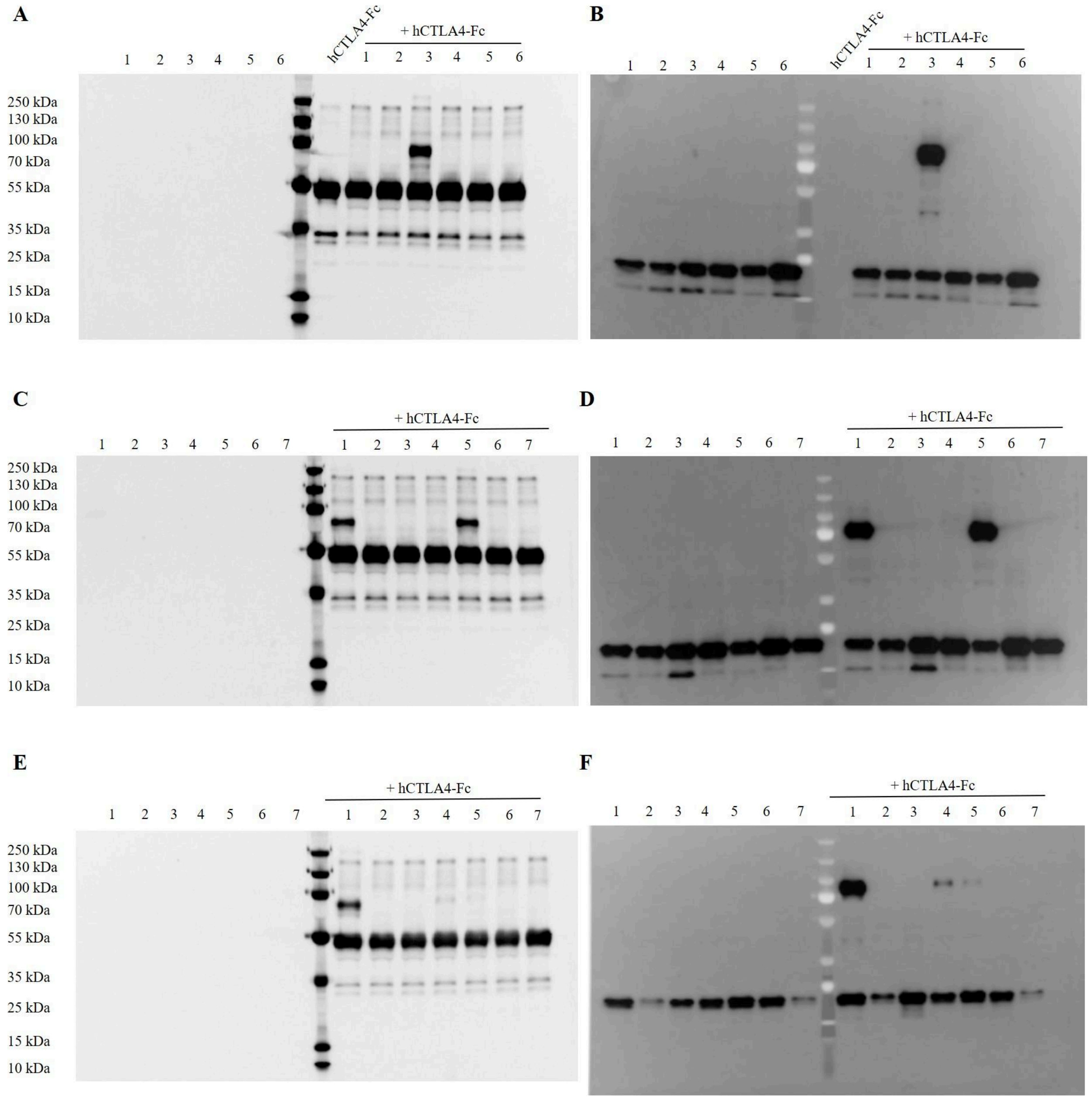


**Fig. S7 | Western blots showing covalent adduct formation between FSY-conjugated 8GAB covalent miniproteins and CTLA-4.** Gels were stained with fluorescent anti-human IgG (A, C, E) and anti-FLAG-HRP (B, D, F). **(A, B)** 1: Y22[FSY], 2: H23[FSY], 3: H24[FSY], 4: H25[FSY], 5:N49[FSY], 6:L50[FSY]. **(C, D)** 1: E51[FSY], 2: Q52[FSY], 3: A53[FSY], 4: V54[FSY], 5: T55[FSY], 6: L56[FSY], 7: V57[FSY]. **(F, E)** 1: S58[FSY], 2: I59[FSY], 3: A60[FSY], 4: S101[FSY], 5: V102[FSY], 6: K103[FSY], 7: M104[FSY].

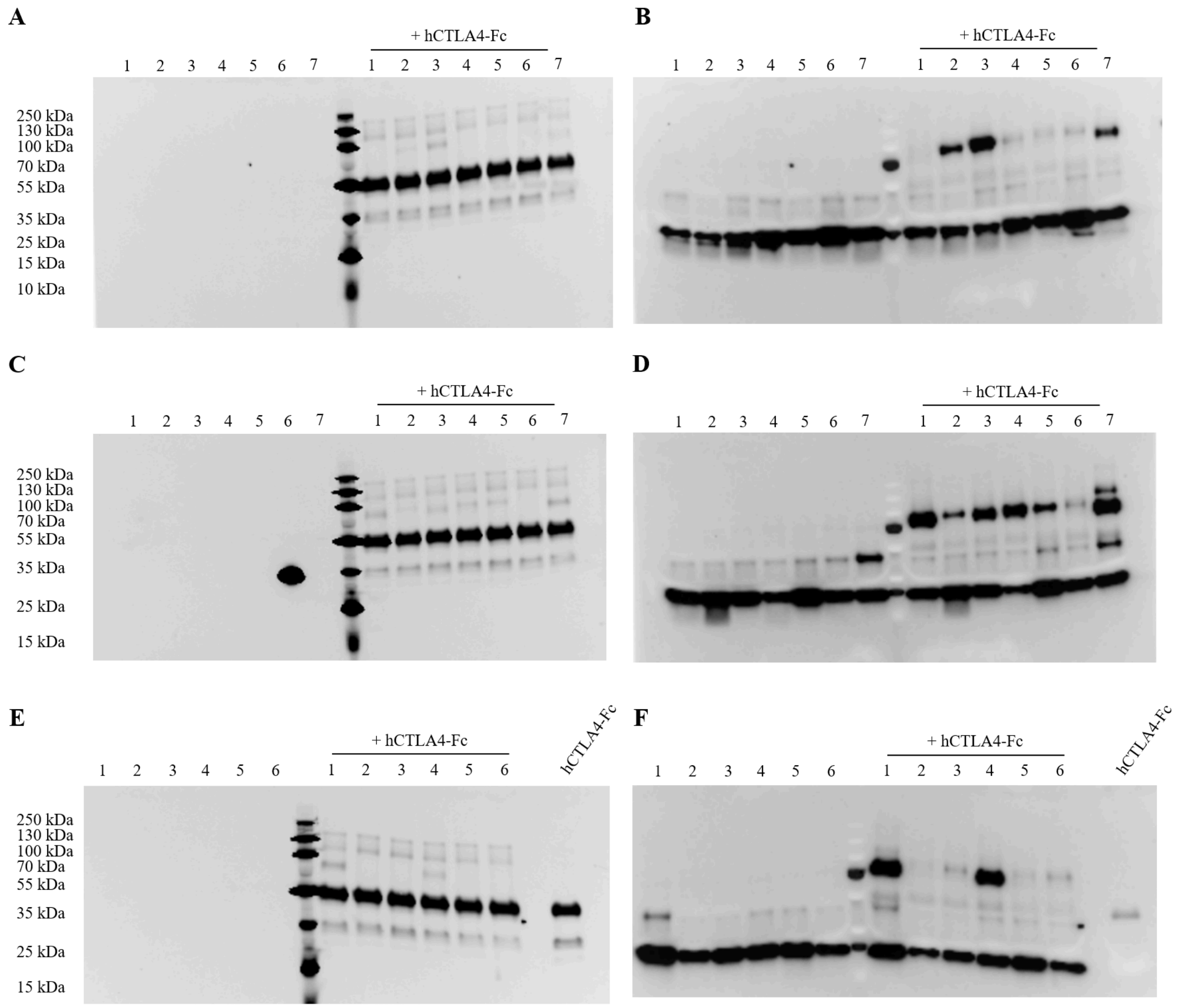


**Fig. S8 | Western blots showing covalent adduct formation between 51C3-conjugated 8GAB covalent miniproteins and CTLA-4.** Gels were stained with fluorescent anti-human IgG (A, C, E) and anti-FLAG-HRP (B, D, F). **(A, B)** 1: Y22[51C3], 2: H23[51C3], 3: H24[51C3], 4: H25[51C3], 5:N49[51C3], 6:L50[51C3], 7:51E[51C3]. **(C, D)** 1: Q52[51C3], 2: A53[51C3], 3: V54[51C3], 4: T55[51C3], 5: L56[51C3], 6: N93[51C3] 7: K94[51C3]. **(F, E)** 1: L95[51C3], 2: A96[51C3], 3: L97[51C3], 4: L98[51C3], 5: V99[51C3], 6: M100[51C3].

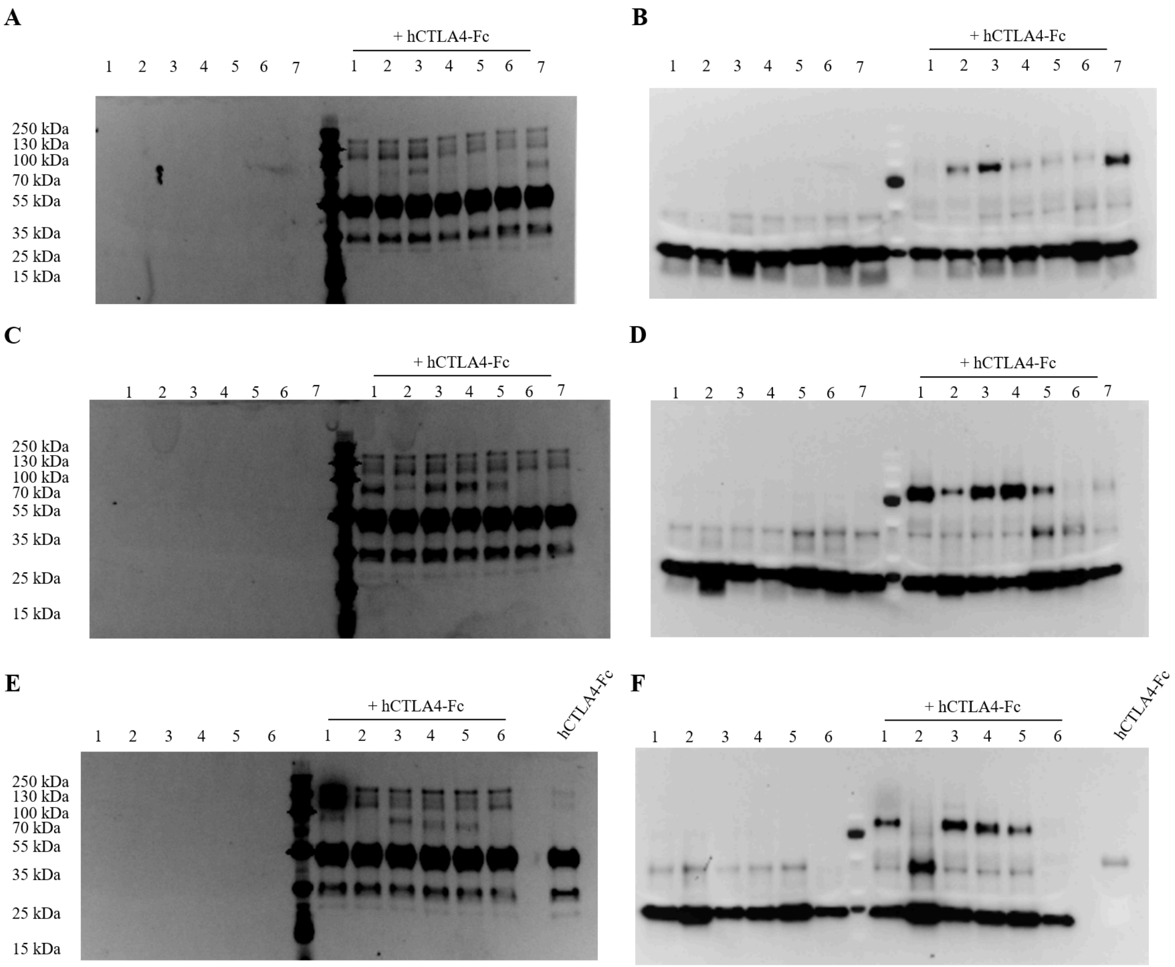


**Fig. S9 | Western blots showing covalent adduct formation between 51C5-conjugated 8GAB covalent miniproteins and CTLA-4.** Gels were stained with fluorescent anti-human IgG (A, C, E) and anti-FLAG-HRP (B, D, F). **(A, B)** 1: Y22[51C5], 2: H23[51C5], 3: H24[51C5], 4: H25[51C5], 5:N49[51C5], 6:L50[51C5], 7:51E[51C5]. **(C, D)** 1: Q52[51C5], 2: A53[51C5], 3: V54[51C5], 4: T55[51C5], 5: L56[51C5], 6: V57[51C5], 7: S58[51C5]. **(F, E)** 1: I59[51C5], 2: A60[51C5], 3: N93[51C5], 4: K94[51C5], 5: L95[51C5], 6: A96[51C5].

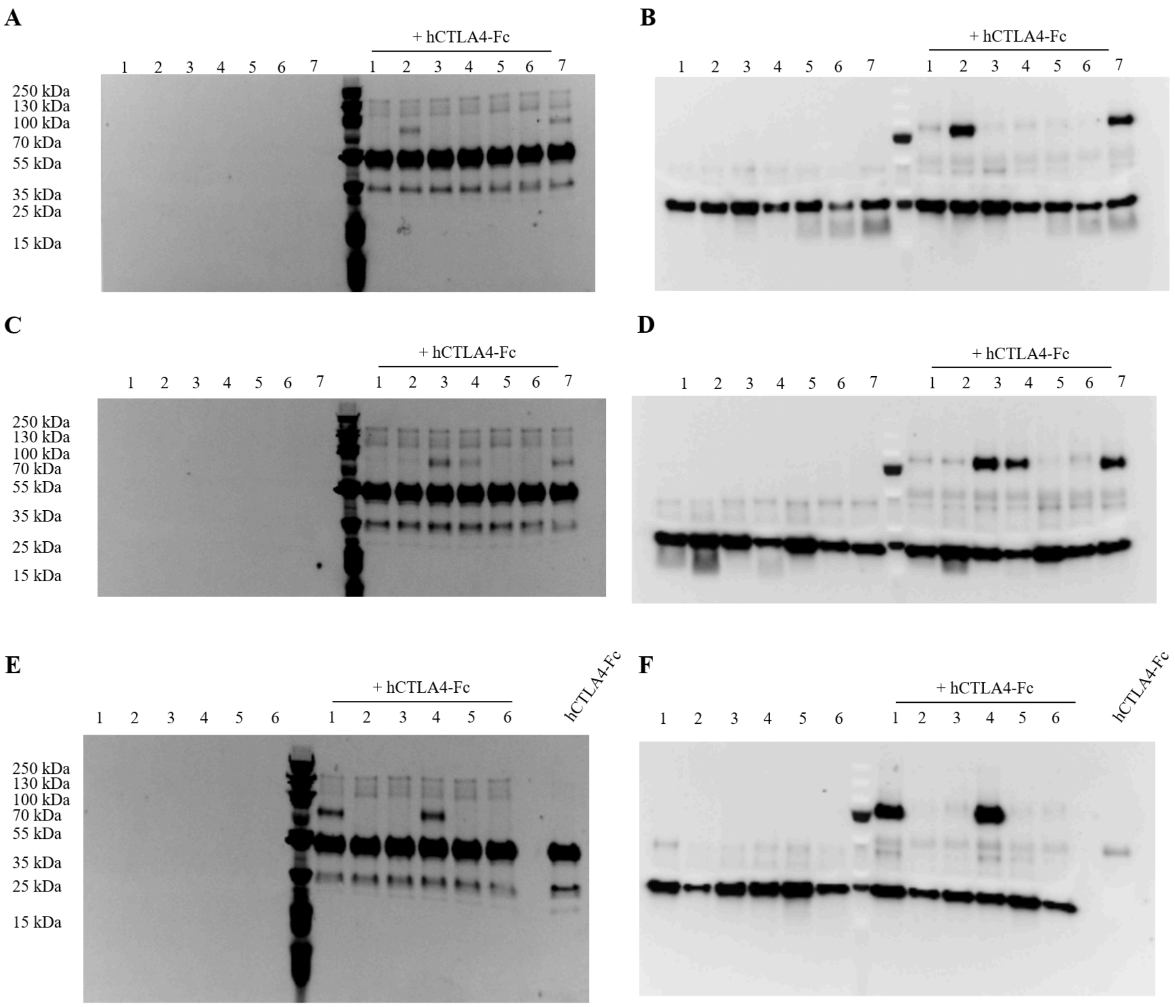


**Fig. S10 | Western blots showing covalent adduct formation between 51C5 and 51C4Me-conjugated 8GAB covalent miniproteins and CTLA-4.** Gels were stained with fluorescent anti-human IgG (A, C, E) and anti-FLAG-HRP (B, D, F). **(A, B)** 1: L97[51C5], 2: L98[51C5], 3: V99[51C5], 4: M100[51C5], 5: Y22[51C4Me], 6: H23[51C4Me], 7: H24[51C4Me]. **(C, D)** 1: H25[51C4Me], 2: A53[51C4Me], 3: V54[51C4Me], 4: T55[51C4Me], 5: L56[51C4Me], 6: N93[51C4Me], 7: K94[51C4Me]. **(F, E)** 1: L95[51C4Me], 2: A96[51C4Me], 3: L97[51C4Me], 4: L98[51C4Me], 5: V99[51C4Me], 6: M100[51C4Me].

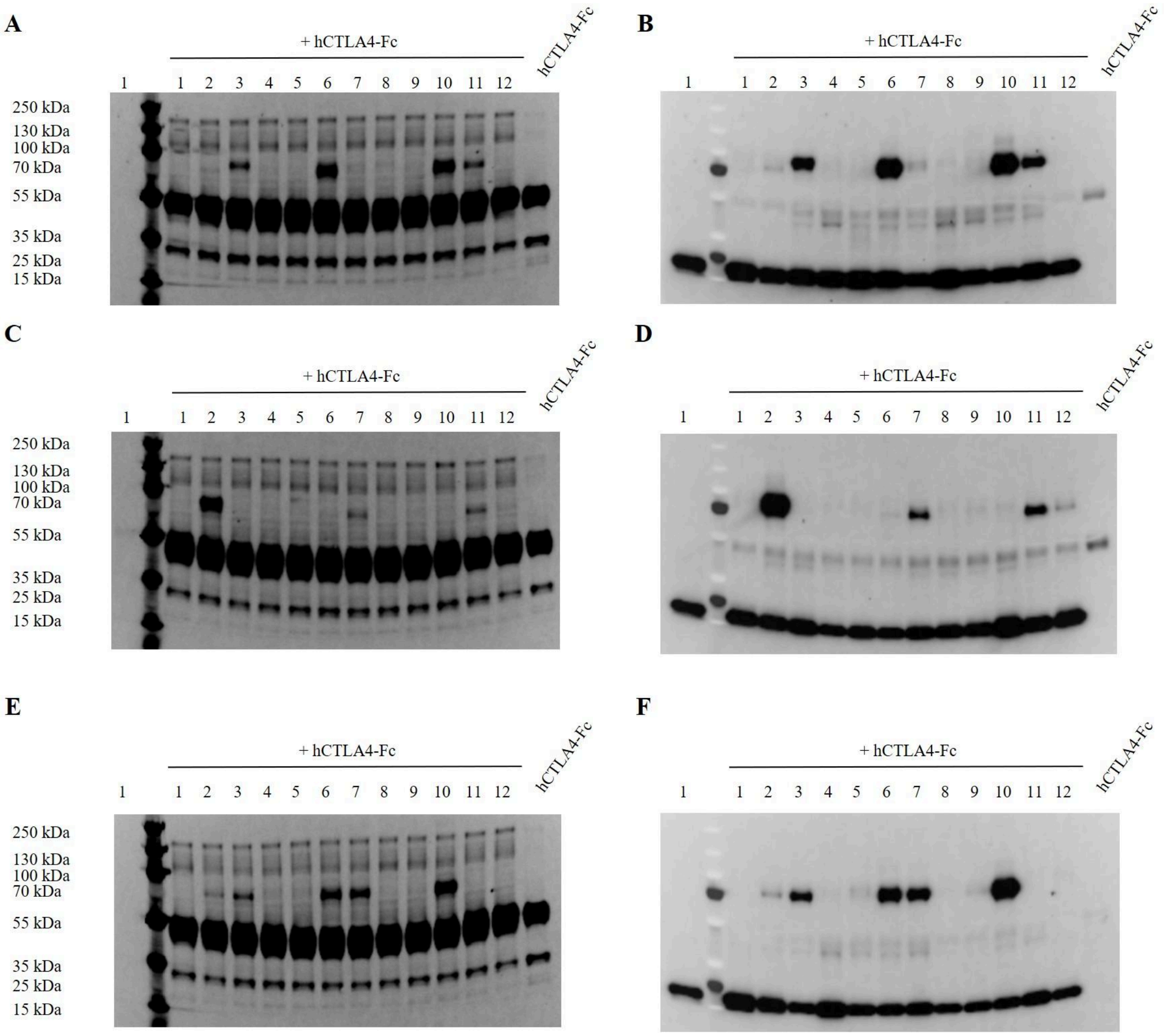


**Fig. S11 | Western blots showing covalent adduct formation between 1C1 and 1C2-conjugated 8GAB covalent miniproteins and CTLA-4.** Gels were stained with fluorescent anti-human IgG (A, C, E) and anti-FLAG-HRP (B, D, F). **(A, B)** 1: Y22[1C1], 2: H23[1C1], 3: H24[1C1], 4: H25[1C1], 5: A53[1C1], 6: V54[1C1], 7: T55[1C1], 8: L56[1C1], 9: N93[1C1], 10: K94[1C1], 11: L95[1C1], 12: A96[1C1]. **(C, D)** 1: L97[1C1], 2: L98[1C1], 3: V99[1C1], 4: M100[1C1]. 5: Y22[1C2], 6: H23[1C2], 7: H24[1C2], 8: H25[1C2], 9: N49[1C2], 10: L50[1C2], 11: 51E[1C1], 12: Q52[1C2]. **(E, F)** 1:A53[1C2], 2: V54[1C2], 3: T55[1C2], 4: L56[1C2], 5:N93[1C2], 6: K94[1C2], 7: L95[1C2], 8: A96[1C2], 9: L97[1C2], 10: L98[1C2], 11: V99[1C2], 12: M100[1C2].

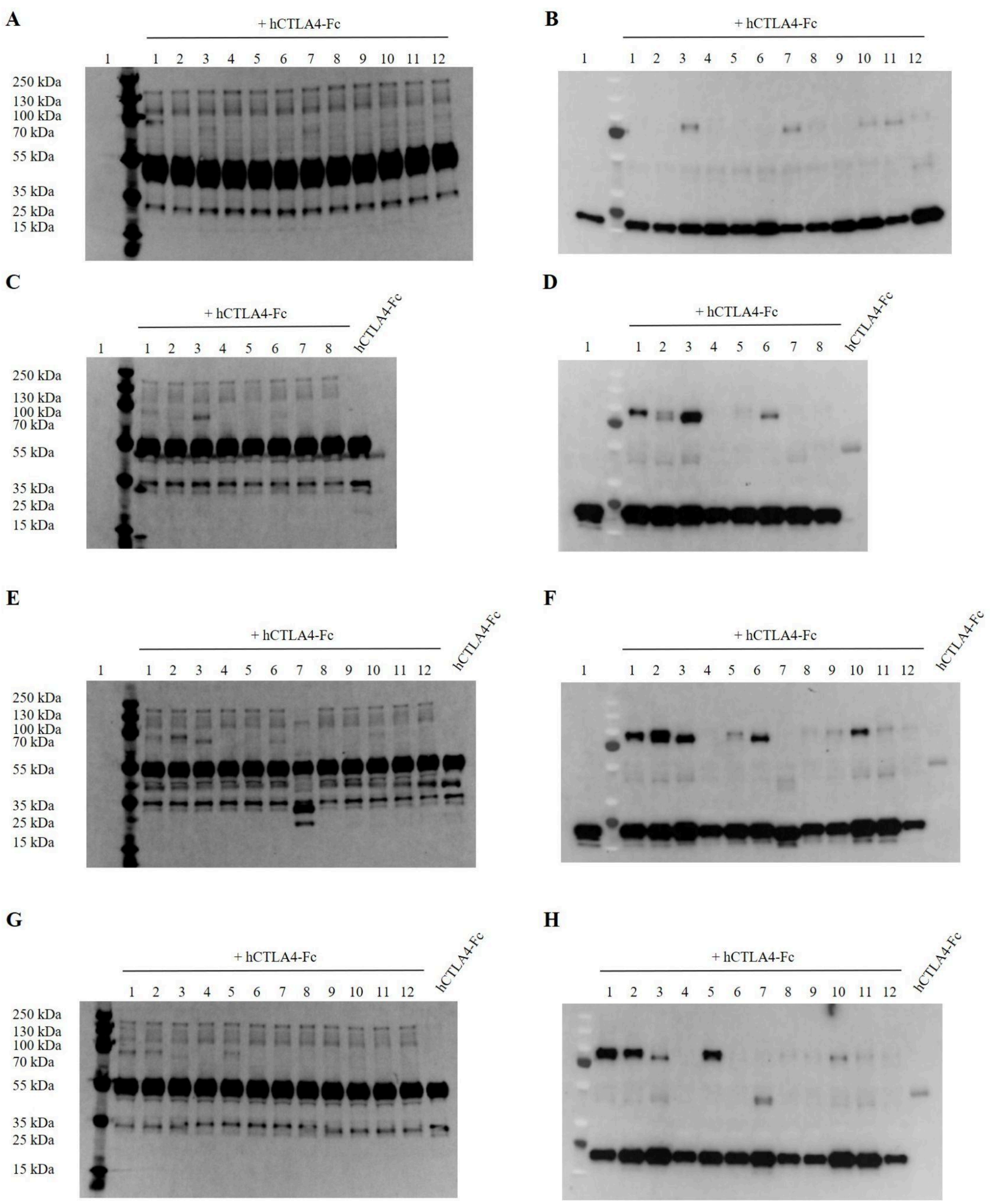


**Fig. S12 | Western blots showing covalent adduct formation between 1C3, 1C4 and 1C5-conjugated 8GAB covalent miniproteins and CTLA-4.** Gels were stained with fluorescent anti-human IgG (A, C, E, G) and anti-FLAG-HRP (B, D, F, H). **(A, B)** 1: Y22[1C3], 2: H23[1C3.], 3: H24[1C3], 4: H25[1C3], 5: N49[1C3], 6: L50[1C3], 7: 51E[1C3], 8: Q52[1C3], 9: A53[1C3], 10: V54[1C3], 11: T55[1C3], 12: L56[1C3]. **(C, D)** 1: N93[1C3], 2: K94[1C3], 3: L95[1C3], 4: A96[1C3], 5: L97[1C3], 6: L98[1C3], 7: V99[1C3], 8: M100[1C3]. **(E, F)** 1: N93[1C4], 2: K94[1C4], 3: L95[1C4], 4: A96[1C4], 5: L97[1C4], 6: L98[1C4], 7: V99[1C4], 8: M100[1C4], 9: S101[1C4], 10: V102[1C4], 11: K103[1C4], 12: M104[1C4]. **(G, H)** 1:

N93[1C5], 2: K94[1C5], 3: L95[1C5], 4: A96[1C5], 5: L97[1C5], 6: L98[1C5], 7: V99[1C5], 8: M100[1C5], 9: S101[1C5], 10: V102[1C5], 11: K103[1C5], 12: M104[1C5].

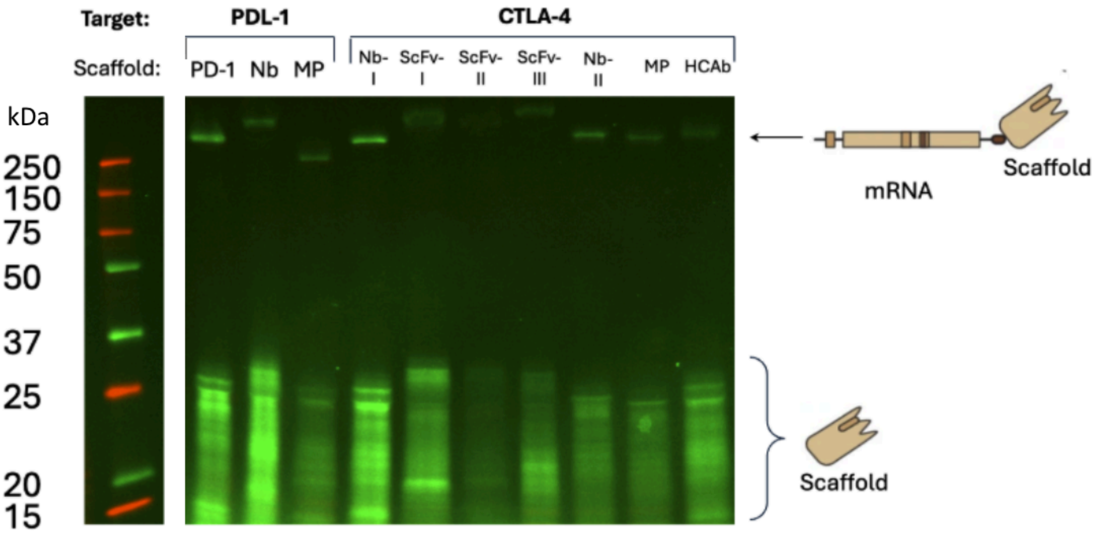


**Fig. S13 | mRNA display library generation across diverse scaffolds and targets**. TAG scan libraries pooled and translated with BDPaF (p-(BODIPY-FL)-aminophenylalanine) via BDPaF-tRNA$_{CUA}$ and displayed as mRNA-protein fusions. Scaffolds include: native receptor-ligand pairs (PD-1 ectodomain targeting PD-L1), anti-PD-L1 nanobody, and multiple anti-CTLA-4 binders including nanobodies (Nb-I: KN44, Nb-II: Nb16), scFvs (ScFv-I: ipilimumab, ScFv-II: JS7, ScFv-III: tremelimumab), miniprotein (MP: 8GAB), and heavy-chain antibody (HCAb 4003-2). Upper band (arrow): mRNA-protein fusions; lower bands (bracket): free translated protein. In-gel fluorescence imaging (ladder in kDa).

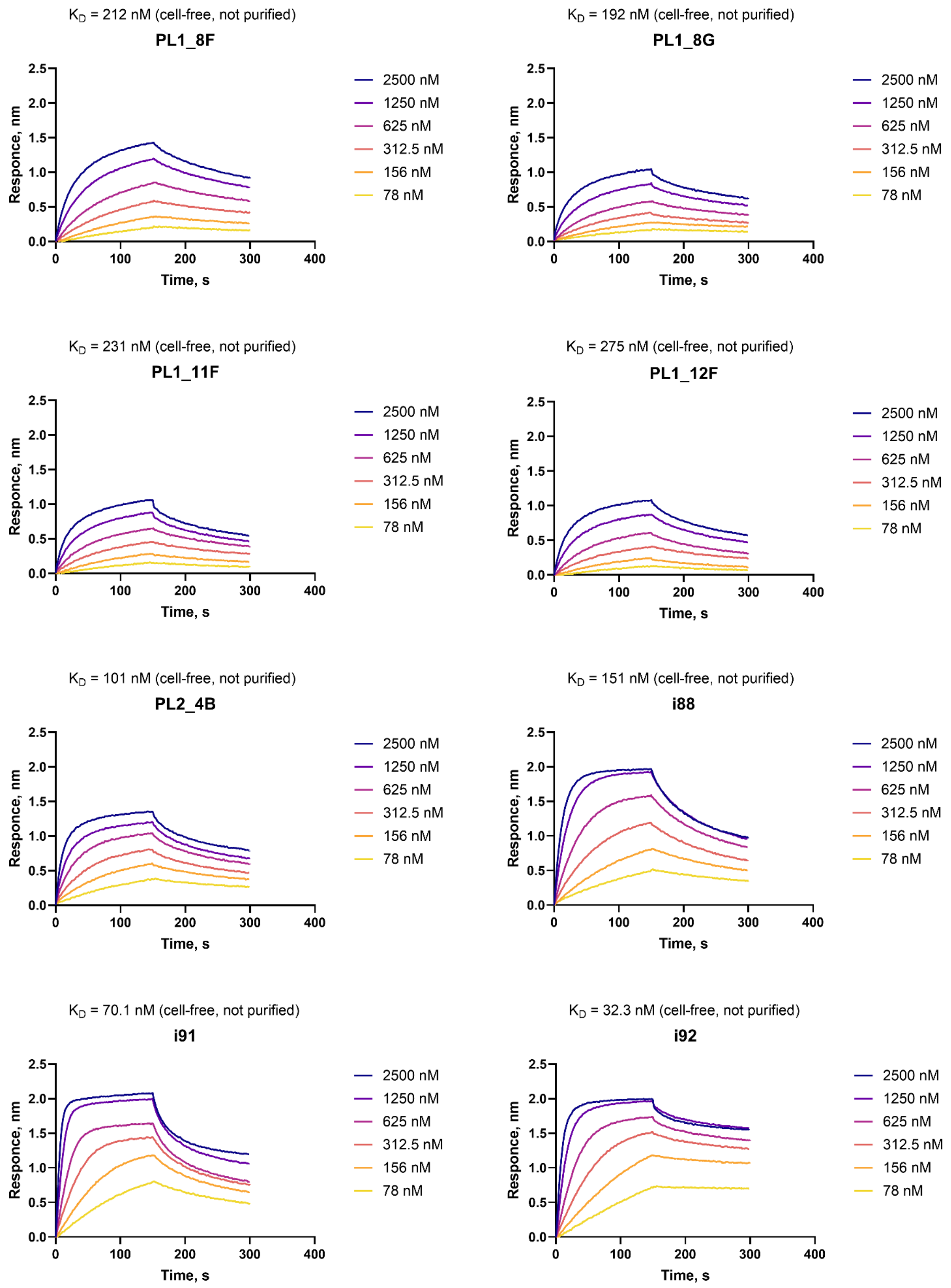


**Fig. S14 | Biolayer interferometry of top-performing de novo miniproteins designed with Bindcraft.** Miniproteins were expressed in cell-free, conjugated on StrepTactin biosensors on a BLI (Gator) and assayed for binding to CTLA-4.

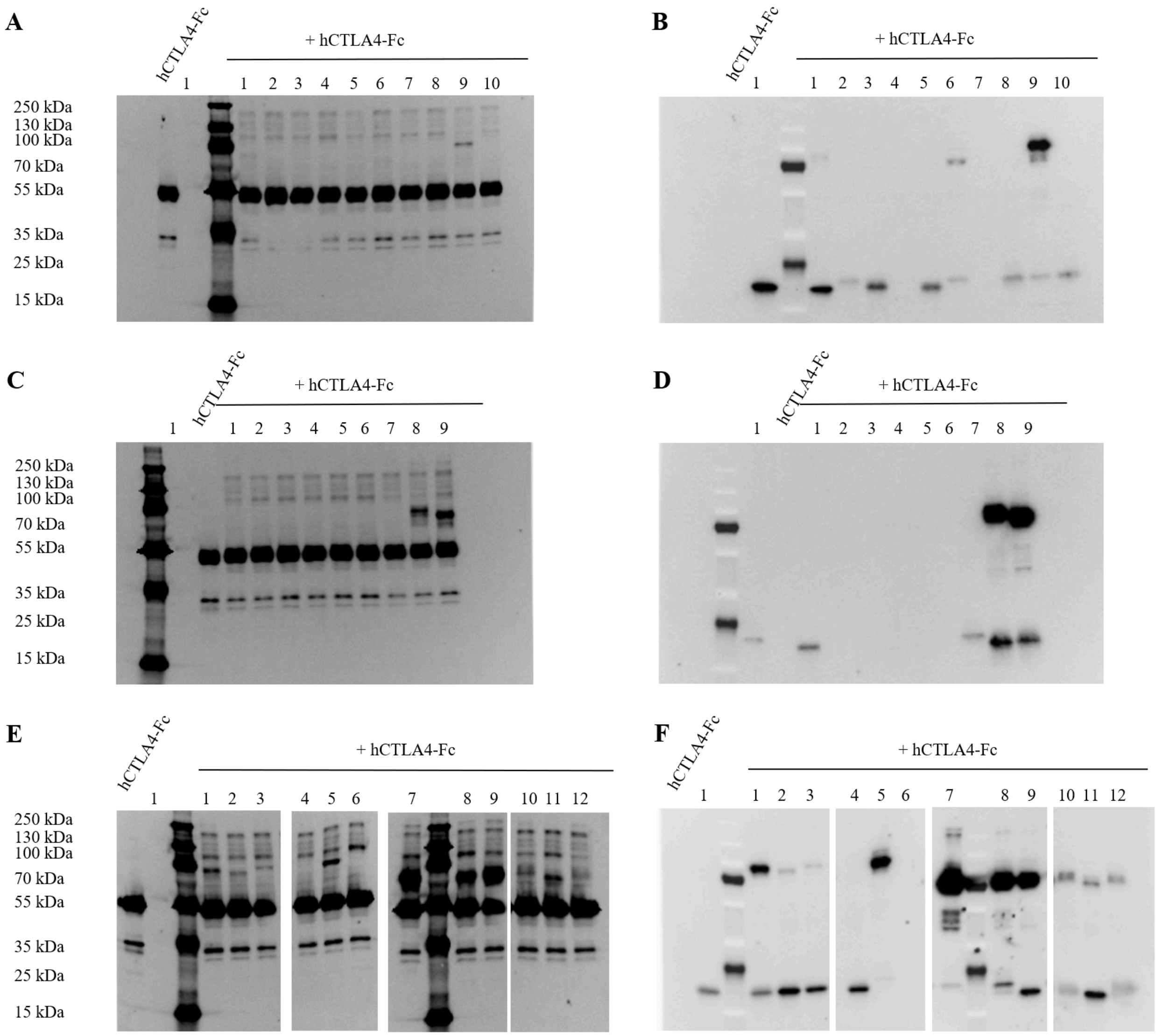


**Fig. S15 | Western blots showing covalent adduct formation between covalent miniproteins and CTLA-4 (part I).** Gels were stained with fluorescent anti-human IgG (A, C, E, G) and anti-FLAG-HRP (B, D, F, H). **(A, B)** 1: I88-44H[1C4], 2: I91-72Q[1C4], 3: I92-45R[1C4], 4: I91-73E[1C4], 5: I92-73E[1C4], 6: I91-58E[1C5], 7: I88-23N[FSY], 8: I91-57V[FSY], 9: I91-44E[FSY], 10: I91-50Y[FSY]. **(C, D)** 1: I92-47H[51C3], 2: I88-22E[51C3], 3: I88-23N[51C5], 4: I92-54R[51C5], 5: I88-22E[51C5], 6: I92-87A[51C5], 7: I91-54L[FSY], 8: I92-63H[FSY], 9: I92-71L[FSY]. **(E, F)** 1: I92-59E[1C4], 2: I88-45T[1C4], 3: I88-68R[1C4], 4: I88-50L[1C5], 5: I91-70Y[FSY], 6: I88-23N[FSY], 7: I91-59I[FSY], 8:I91-55H[FSY], 9: I88-51E[FSY], 10; I92-88E[FSY], 11: I88-47E[FSY], 12: I92-89F[FSY].

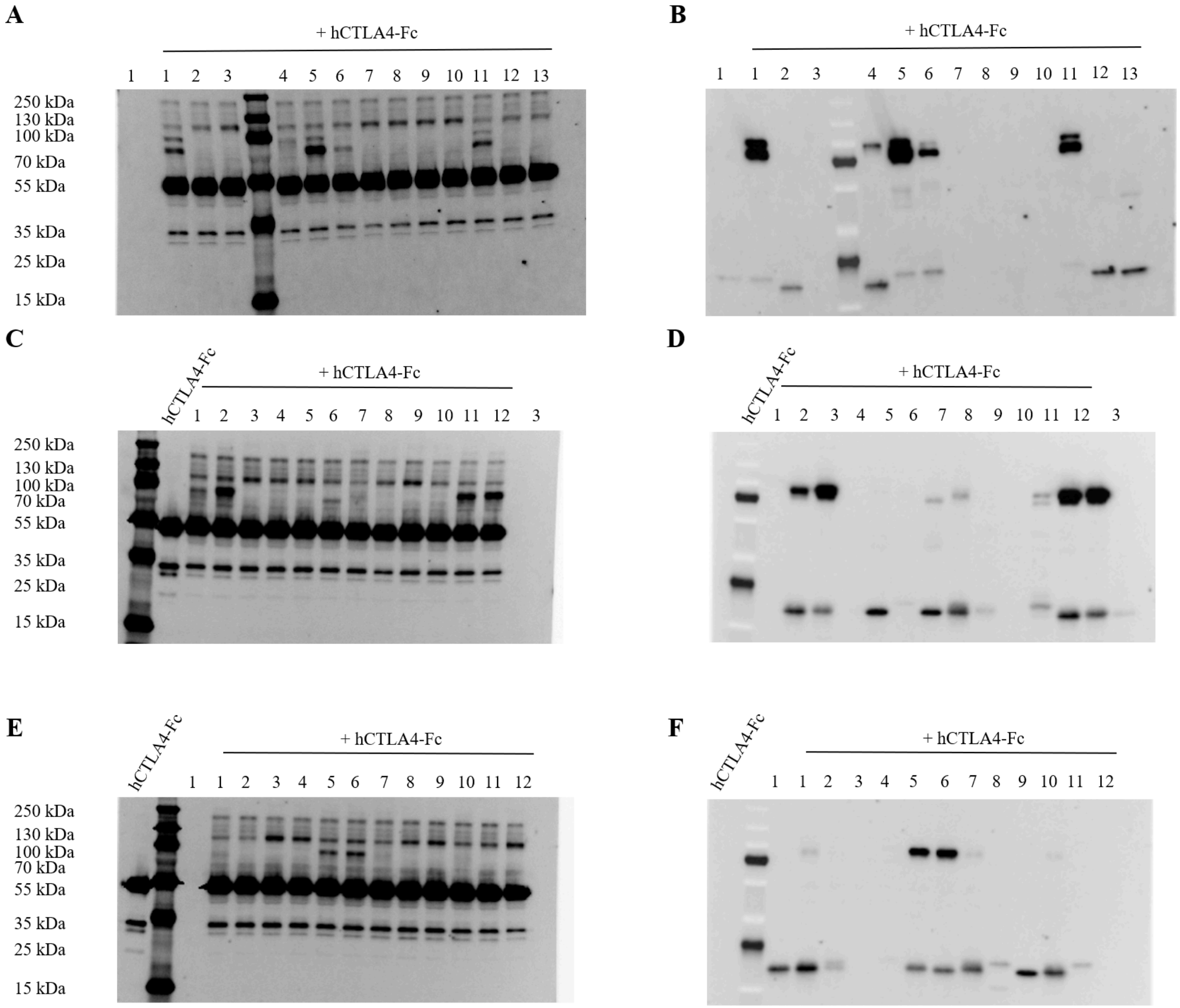


**Fig. S16 | Western blots showing covalent adduct formation between *de novo* miniproteins and CTLA-4 (part II).** Gels were stained with fluorescent anti-human IgG (A, C, E, G) and anti-FLAG-HRP (B, D, F, H). **(A, B)** 1: I91-58E[51C3], 2: I92-47H[51C3], 3: I88-22E[51C3], 4: I92-59E[51C4Me], 5: I91-58E[51C4Me], 6: I91-64S[51C4Me], 7: I88-23N[51C5] 8: I92-54R[51C5], 9: I88-22E[51C5], 10: I92-87A[51C5], 11: I91-54L[51C5], 12: I88-28C[51C5], 13: I88-25R[51C5]. **(C, D)** 1: I92-59E[1C1], 2: I92-57P[1C1], 3: I88-23N[1C1], 4: I88-68R[1C1], 5: I91-47E[1C1], 6: I88-58G[1C1], 7: I92-63H[1C1], 8: I92-41C[1C1], 9: I92-70A[1C1], 10: I91-64S[1C2], 11: I92-59E[1C2], 12: I92-57P[1C2]. **(E, F)** 1: I92-66E[1C2], 2: I91-89E[1C2], 3: I92-70A[1C2], 4: I91-47E[1C3], 5: I92-57P[1C3], 6: I92-59E[1C3], 7: I92-76A[1C3], 8: I91-42R[1C3], 9: I88-67L[1C3], 10: I92-77L[1C3], 11: I91-79A[1C3], 12: I88-29R[1C3].

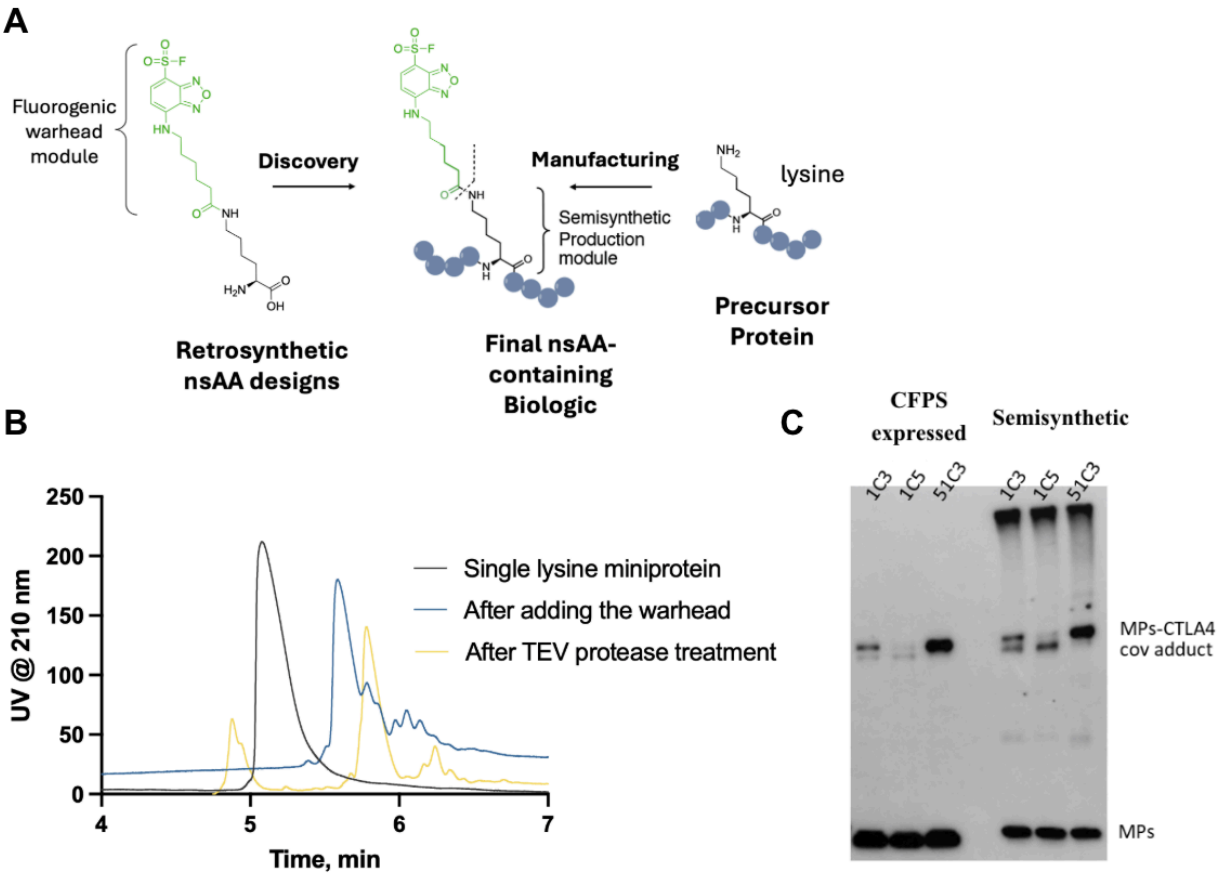


**Fig. S17 | Retrosynthetic nsAA design enables scalable semisynthetic production. A.** (Left) Lysine-based non-standard amino acid (nsAA) with modular warhead (51C6, green) used for mRNA display screening is reformatted as an NHS ester and conjugated post-translationally to a precursor protein containing a single lysine residue at the identified position. (Right) Semisynthetic conjugation of NHS-activated warheads to single-lysine miniproteins. **B.** Representative HPLC chromatogram for 8GAB with 51C4-Me at position 51 showing the semisynthetic conjugation of NHS-activated warheads to single-lysine miniproteins. Lysine ε-amino group-specific modification of the miniprotein is confirmed by incorporating a TEV protease-cleavable sequence at the N-terminus and subsequent treatment with TEV protease, which separates the warhead-conjugated miniprotein from the unmodified N-terminal peptide fragment. **C.** Western blot comparison of covalent CTLA-4 miniprotein 8GAB with no lysine variants produced via PURExpress cell-free translation (CFPS expressed) or two-step chemistry (semisynthetic, recombinant expression followed by NHS-ester conjugation). Miniproteins were modified with 1C3, 1C5, or 51C3 warheads. StrepTactin-HRP detection shows miniproteins (MPs, lower bands) and covalent miniprotein-CTLA-4 adducts (MPs-CTLA-4 covalent adduct, upper bands), demonstrating comparable covalent reactivity of miniproteins produced by CFPS and scaled-up semisynthetically.

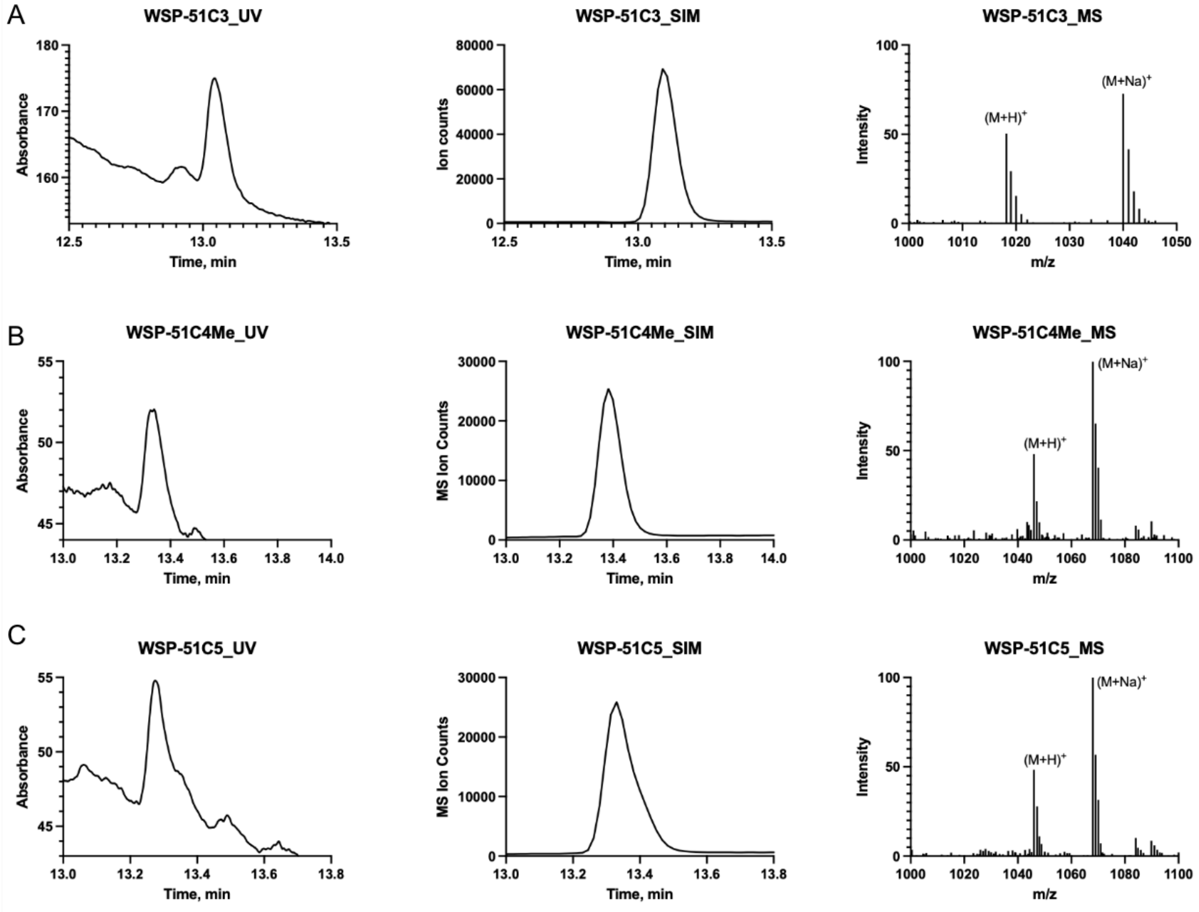


**Fig. S18 | LC-MS verification of site-specific ribosomal incorporation of novel fluorogenic amino acids.** Fluorogenic amino acids with varying side chain lengths (WSP-51C3, WSP-51C4-Me, and WSP-51C5) were incorporated at position * of the hexapeptide fMFPV*V. UV absorbance at 210 nm (left panel), extracted ion chromatograms (middle panel), and corresponding mass spectra (right panel) for fMFPV*V peptides containing the non-standard amino acids: (A) WSP-51C3, (B) WSP-51C4Me, and (C) WSP-51C5.

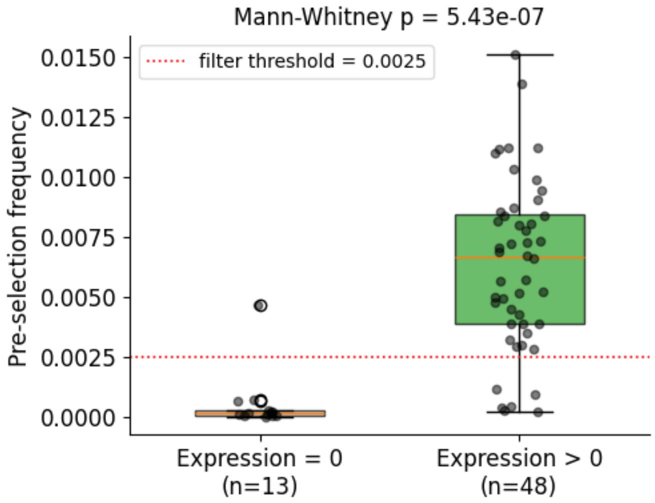


**Fig. S19 | Pre-selection frequency stratified by expression for the ML-designed miniproteins.** Box plots compare the pre-selection frequency of variants that failed to express in the western blot assay (Expression = 0, grey) with that of variants that expressed (Expression > 0, green); individual variants are overlaid as black dots. The red dotted line indicates the pre-selection frequency threshold (0.0025) applied to filter rare reads in downstream analyses. The two distributions differ significantly (Mann-Whitney U test, two-sided), confirming that variants with very low pre-selection frequency are disproportionately non-expressing and motivating the threshold choice.

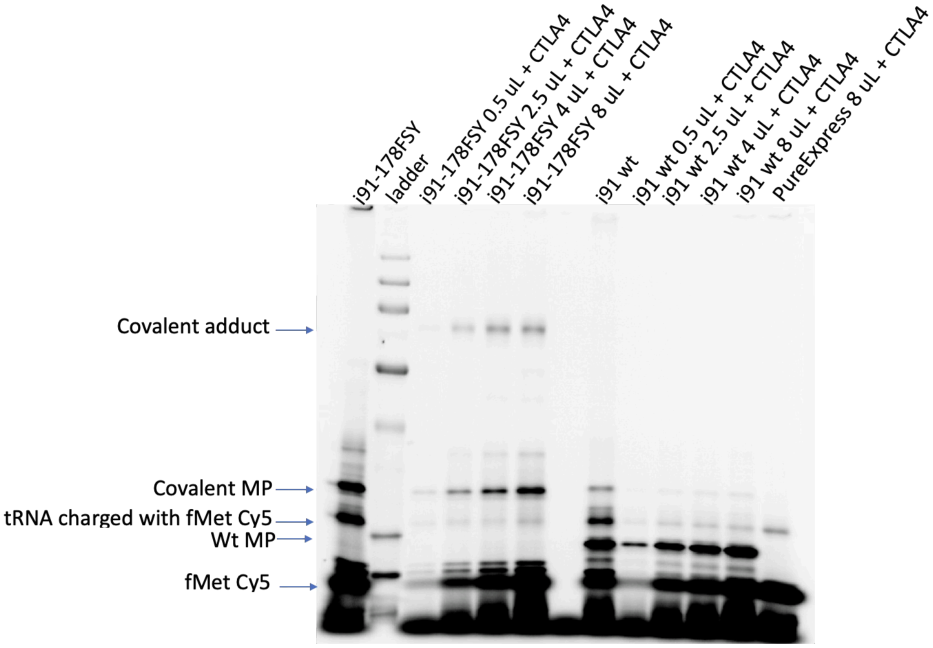


**Fig. S20 | Competitive dissociation assay normalized by N-terminal fluorescent labeling.** Covalent (I91 59[FSY]) and non-covalent (I91 wt) variants were expressed with Cy5-Met at the N-terminus via misaminoacylated initiator tRNA. Because Cy5-Met incorporation reflects relative nascent protein synthesis, different volumes of cell-free reactions were loaded to achieve equal protein amounts based on

Cy5 fluorescence intensity. Normalized proteins were then mixed at defined ratios with CTLA-4 and incubated to allow covalent bond formation. Gel electrophoresis with in-gel fluorescence detection shows relative protein loading (bottom bands) and formation of covalent adduct (upper bands) only for the FSY-containing variant. Mobility differences between variants reflect different affinity tags (StrepTactin for wt, FLAG for TAG scan variant). The same Cy5-Met label enables both expression normalization and direct visualization of covalent complex formation.

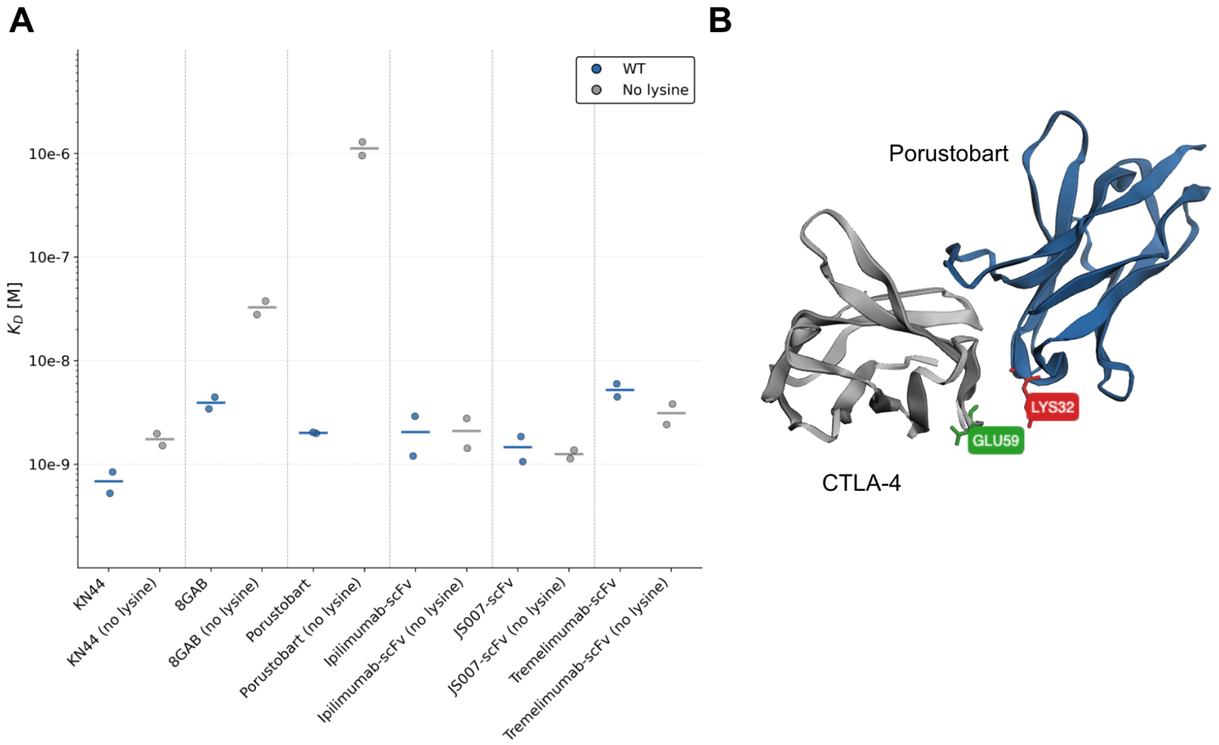


**Fig. S21 | Global lysine-to-arginine substitution preserves binding affinity across diverse scaffold types. A.** Binding affinities ($K_d$) of CTLA-4 binders were measured by BLI and compared between wild-type (WT) scaffolds and their lysine-less (no lysine, with K→R substitutions) variants. Miniproteins (8GAB), scFvs (ipilimumab-scFv, tremelimumab-scFv, JS007-scFv) and nanobodies (KN44) retained binding affinities in the nanomolar range after complete lysine substitution with arginine. Lines represent mean $K_d$ values with individual replicates shown as dots. **B.** Crystal structure of porustobart bound to CTLA-4 (PDB: 7DV4) [4]. The no-lysine version showed reduced affinity likely due to K32 in the binding interface forming a salt bridge with CTLA-4 E59.

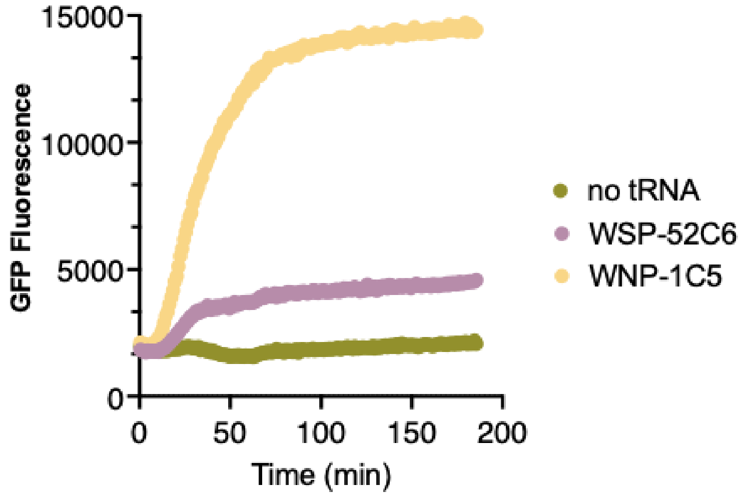


**Fig. S22 |** GFP reporter assay for fluorogenic nsAA incorporation. Engineered sfGFP templates contain a TAG (amber) codon at position 2. Successful incorporation of nsAAs (WNP-1C5, WSP-52C6) during translation produces GFP, while no-tRNA controls show minimal background fluorescence.

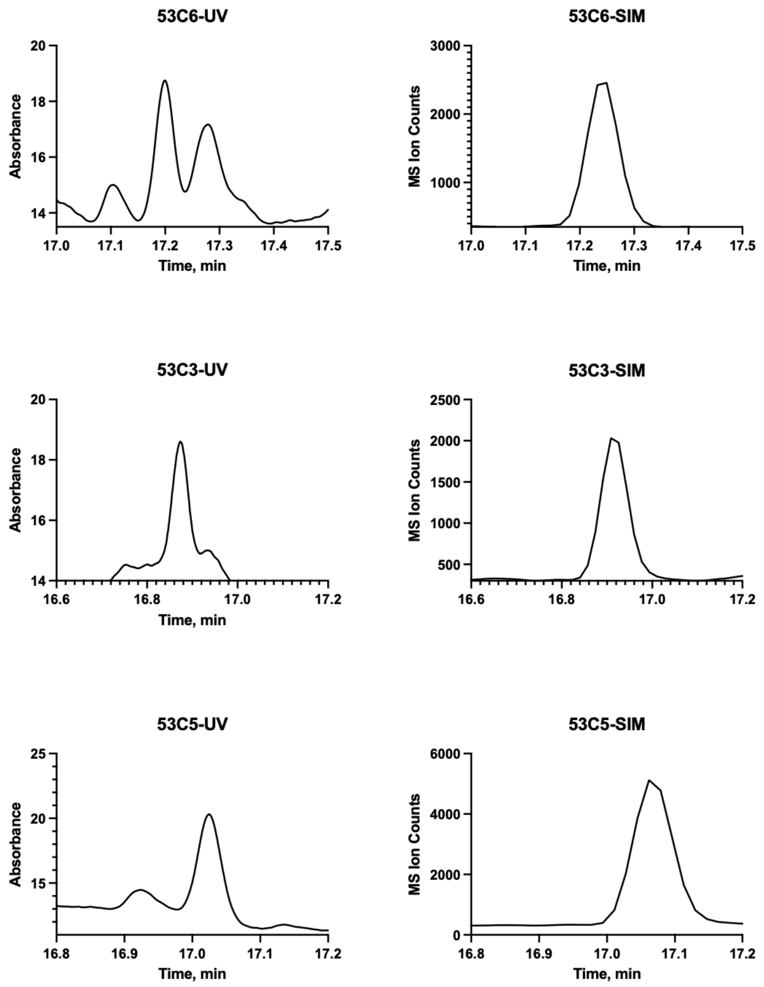


**Fig. S23 | HPLC-MS verification of site-specific ribosomal incorporation of fluorogenic amino acids.** Fluorogenic amino acids with varying side chain lengths (WSP-53C3, WSP-53C5, and WSP-53C6) were incorporated at position * of the hexapeptide fMFPV*V. UV absorbance at 210 nm (left panel), extracted ion chromatograms (right panel), confirm site-specific incorporation.

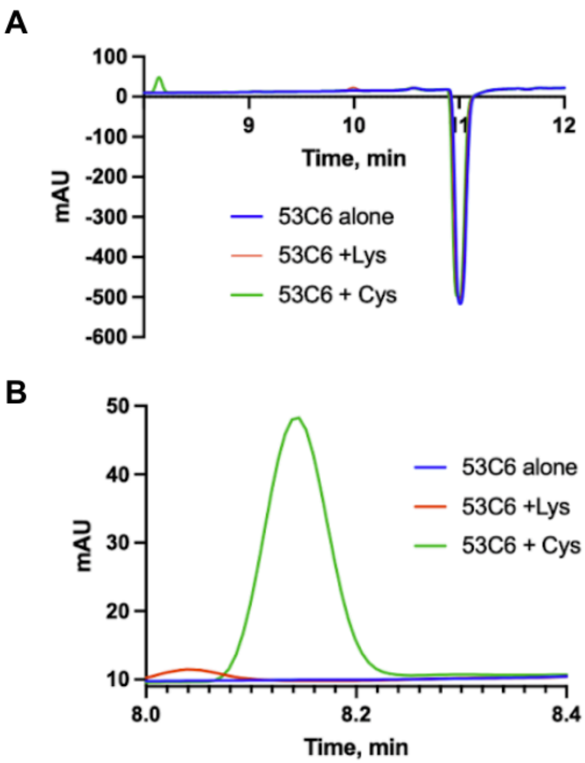


**Fig. S24 | HPLC analysis of 53C6 covalent reactivity**. A. Chromatograms show 53C6 (blue) reaction with target nucleophiles (cysteine in green/lysine in red), demonstrating warhead reactivity and covalent bond formation. **B.** Zoom on the left peak in panel A.

### Supplementary Tables

**Table S1. Key DNA or RNA sequences used in this study.**

| Name | Sequence |
|---|---|
| 8GAB-S1TAG | CTAGCCCCGCGAAATTAATACGACTCACTATAGGGTCTAGA AATAATTTTGTTTAACTTTAAGAAGGAGATATACATATGAAA ATCGAAGAATCAGGGCCGAGTCGCCTGGAAGAGGAATTAC GCCGCCGCCTGACAGAACCGTCGGGTTAGGGCTCCGGTAA CAACTTAGAGGAAGAGGTCCACAAGCTGCTGTGGGAACTG TCTGAAATTTACCACCACCATGACCATGAAGCATCCCATGA GGCGCTGCGCCGCGCCCTGGAAGTGCTGAAGCAGTTGCTT GAACACAACAATTTAGAACAAGCTGTGACTTTAGTTTCGAT TGCTGTGCATGTAGCGGTCCGTGTCAATGACGAGCATGTAA TCCGTGAATTGCGTCACTTCCTTCGCCGTTTACTGAAACAG |

| | |
|---|---|
| | GTAAAGGAGCACAATAACAATAAGCTGGCTTTGCTTGTGAT GTCAGTAAAAATGCAGTTGGATCGCACCAGCGGCGGGGGA GGTTCAGATTACAAGGATCACGATGGAGACTATAAGGATCA CGACATCGATTACAAAGACGATGACGACAAAGGCGGTGGA GGCTCAGGTGGGGGATCTGGTGGTCACCATCACCACCATC ATGGAAGCGGCCATCACCATCATCATCATGGTTCTGGTGGT GGTTCTGGTTAATGAGGATCCCGGGAATTCTCGAGTAAGGT TAACCTG |
| I88-D21TAG | CTAGCCCCGCGAAATTAATACGACTCACTATAGGGTCTAG AAATAATTTTGTTTAACTTTAAGAAGGAGATATACATATGA AAATCGAAGAATCAGGGCCGAGTCGCCTGGAAGAGGAA TTACGCCGCCGCCTGACAGAACCGTCGGGTAGCTGGCGC GAAGTGGTGGAACGCTTTCGCGAAATTGCGGAAGAAAC CGCGGAACGCTTTCCGTAGGAAAACGTGCGCTTTTGCTG CCGCATTGTGGTGCAGGATTGCGATCATATGATGGAATAT GGCGGCCATACCGAAGAACAGACCCTGGAAGCGGGCCG CGCGAGCATTGGCCTGGCGATTCGCCTGCTGAGCGGCCT GCGCGGCAACGAAGCGGCGGCGCGCGCGGTGGAATATCT GCGCGCGCGCCTGGCGGAACTGAGCGGCGGGGGAGGTT CAGATTACAAGGATCACGATGGAGACTATAAGGATCACG ACATCGATTACAAAGACGATGACGACAAAGGCGGTGGAG GCTCAGGTGGGGGATCTGGTGGTCACCATCACCACCATC ATGGAAGCGGCCATCACCATCATCATCATGGTTCTGGTGG TGGTTCTGGTTAATGAGGATCCCGGGAATTCTCGAGTAAG GTTAACCTG |
| I91-R42TAG | CTAGCCCCGCGAAATTAATACGACTCACTATAGGGTCTAG AAATAATTTTGTTTAACTTTAAGAAGGAGATATACATATGA AAATCGAAGAATCAGGGCCGAGTCGCCTGGAAGAGGAA TTACGCCGCCGCCTGACAGAACCGTCGGGTAGCGAAGAA GAACGCCGCATTGAATGGGGCCGCCGCATGGTGGAACTG ATGCGCGAAGCGGCGGAAAACCCGGAACGCATGGAAGA ACTGGCGGAAGAAGTGCGCCGCCTGAGCGAAGGCATGC CGTAGACCGAAGAATTTGAATATCCGTATTGGTATTTTCTG CATTATGTGGAAATTTATCCGAGCCTGAGCGAACCGATGC AGCGCTATATGCAGGAAGAACTGCTGCGCCGCGCGGATG AACTGGAAGAACGCGTGCCGGCGGAACTGAGCGGCGGG GGAGGTTCAGATTACAAGGATCACGATGGAGACTATAAG GATCACGACATCGATTACAAAGACGATGACGACAAAGGC GGTGGAGGCTCAGGTGGGGGATCTGGTGGTCACCATCAC CACCATCATGGAAGCGGCCATCACCATCATCATCATGGTT |

| | |
|---|---|
| | CTGGTGGTGGTTCTGGTTAATGAGGATCCCGGGAATTCTCGAGTAAGGTTAACCTG |
| I92-G31TAG | CTAGCCCCGCGAAATTAATACGACTCACTATAGGGTCTAGAAATAATTTTGTTTAACTTTAAGAAGGAGATATACATATGAAAATCGAAGAATCAGGGCCGAGTCGCCTGGAAGAGGAATTACGCCGCCGCCTGACAGAACCGTCGGGTAGCCGCGAAGAAGTGCGCGAACTGGTGGAACGCGTGCGCACCGGCGATCGCGAAGAACGCCTGGAAGTGCTGGAAGAACTGGAAGAACTGGCGTAGCCGGAATTTCCGATGGCGCGCTATCTGTGCCGCGATCTGCGCCGCCATCTGGAAGAAATGGAAGAACGCAACGAACCGCAGGAAAACGTGGATCATATGATTGAACAGCATATGGCGCTGATTGAACCGGCGCTGCTGCAGGGCGAAGGCATTCGCGAAGGCCGCGCGGAATTTGTGTGGAGCGGCGGGGGAGGTTCAGATTACAAGGATCACGATGGAGACTATAAGGATCACGACATCGATTACAAAGACGATGACGACAAAGGCGGTGGAGGCTCAGGTGGGGGATCTGGTGGTCACCATCACCACCATCATGGAAGCGGCCATCACCATCATCATCATGGTTCTGGTGGTGGTTCTGGTTAATGAGGATCCCGGGAATTCTCGAGTAAGGTTAACCTG |
| Pri1 | GTTTAACTTTAAGAAGGAGATATACATATGAAAATCGAAGAATCAGGGCCGAGTCG |
| Pri2 | CTTACTCGAGAATTCCCGGGATCCTCATTAACCAGAACCACCACCAGAACCATGATGATG |
| Pri5 | TCCCTACACGACGCTCTTCCGATCTCCGCCGCCTGACAGAAC |
| Pri6 | GTTCAGACGTGTGCTCTTCCGATCTGTAATCTGAACCTCCCCCGC |
| P7-Universal reverse (upstream of barcode) | GTTAACCTTACTCGAGAATTCCCGGGATCC |
| P3-I92 forward | GCTGGAAGAACTGGAAGAACTGGCG |
| P4-I88 forward | TCCGGATGAAAACGTGCGCTTTTG |
| P5-I91 forward | GCATGGAAGAACTGGCGGAAGAAG |
| P6-8GAB forward | GCGCCCTGGAAGTGCTGAAG |
| R1-Barc-control | GAGAATTCCCGGGATCCTCAC**TTAT**TTTTTTTTTTTTTTTTTTGCACCAGAACCACCGAG |

| | |
|---|---|
| R2-Barc-1C1 | GAGAATTCCCGGGATCCTCAC**CTACGTAT**TATTTTTTTTT TTTTTTTTTGCACCAGAACCACCGAG |
| R3-Barc-1C2 | GAGAATTCCCGGGATCCTCAC**CAGTACGT**TATTTTTTTTT TTTTTTTTTGCACCAGAACCACCGAG |
| R4-Barc-1C3 | GAGAATTCCCGGGATCCTCAC**GTGACTAT**TATTTTTTTTT TTTTTTTTTGCACCAGAACCACCGAG |
| R5-Barc-1C4 | GAGAATTCCCGGGATCCTCAC**ATCGTCGT**TATTTTTTTTT TTTTTTTTTGCACCAGAACCACCGAG |
| R6-Barc-1C5 | GAGAATTCCCGGGATCCTCAC**GCTCATAT**TATTTTTTTTT TTTTTTTTTGCACCAGAACCACCGAG |
| R7-Barc-51C3 | GAGAATTCCCGGGATCCTCAC**GAGCATCT**TATTTTTTTTT TTTTTTTTTGCACCAGAACCACCGAG |
| R8-Barc-51C4Me | GAGAATTCCCGGGATCCTCAC**AGACTCGT**TATTTTTTTTT TTTTTTTTTGCACCAGAACCACCGAG |
| R9-Barc-51C5 | GAGAATTCCCGGGATCCTCAC**TCGCTGAT**TATTTTTTTTT TTTTTTTTTGCACCAGAACCACCGAG |
| R10-Barc-FSY | GAGAATTCCCGGGATCCTCAC**CGATCTGT**TATTTTTTTTT TTTTTTTTTGCACCAGAACCACCGAG |

| | |
|---|---|
| pPURExpress | GCTAGTGGTGCTAGCCCCGCGAAATTAATACGACTCACTA TAGGGTCTAGAAATAATTTTGTTTAACTTTAAGAAGGAGA TATACATATAATGAGGATCCCGGGAATTCTCGAGTAAGGTT AACCTGCAGGAGGCCTTTAATTAAGGTGGTGCGGCCGCG CTAGCGGTCCCGGGGGATCGATCCGGCTGCTAACAAAGC CCGAAAGGAAGCTGAGTTGGCTGCTGCCACCGCTGAGC AATAACTAGCATAACCCCTTGGGGCCTCTAAACGGGTCTT GAGGGGTTTTTTGCTGAAAGGAGGAACTATATCCGGAAG CTTGGCACTGGCCGACCGGGGTCGAGCACTGACTCGCTG CGCTCGGTCGTTCGGCTGCGGCGAGCGGTATCAGCTCAC TCAAAGGCGGTAATACGGTTATCCACAGAATCAGGGGATA ACGCAGGAAAGAACATGTGAGCAAAAGGCCAGCAAAAG GCCAGGAACCGTAAAAAGGCCGCGTTGCTGGCGTTTTTC CATAGGCTCCGCCCCCCTGACGAGCATCACAAAAATCGA CGCTCAAGTCAGAGGTGGCGAAACCCGACAGGACTATAA AGATACCAGGCGTTTCCCCCTGGAAGCTCCCTCGTGCGCT CTCCTGTTCCGACCCTGCCGCTTACCGGATACCTGTCCGC CTTTCTCCCTTCGGGAAGCGTGGCGCTTTCTCATAGCTCA |

CGCTGTAGGTATCTCAGTTCGGTGTAGGTCGTTCGCTCCA
AGCTGGGCTGTGTGCACGAACCCCCCGTTCAGCCCGACC
GCTGCGCCTTATCCGGTAACTATCGTCTTGAGTCCAACCC
GCTAAGACACGACTTATCGCCACTGGCAGCAGCCACTGG
TAACAGGATTAGCAGAGCGAGGTATGTAGGCGGTGCTAC
AGAGTTCTTGAAGTGGTGGCCTAACTACGGCTACACTAG
AAGAACAGTATTTGGTATCTGCGCTCTGCTGAAGCCAGTT
ACCTTCGGAAAAAGAGTTGGTAGCTCTTGATCCGGCAAA
CAAACCACCGCTGGTAGCGGTGGTTTTTTTGTTTGCAAGC
AGCAGATTACGCGCAGAAAAAAAGGATCTCAAGAAGATC
CTTTGATCTTTTCTACGGGGTCTGACGCTCAGTGGAACGA
AAACTCACAGATCCGGGATTTTGGTCATGAGATTATCAAA
AAGGATCTTCACCTAGATCCTTTTAAATTAAAAATGAAGT
TTTAAATCAATCTAAAGTATATATGAGTAAACTTGGTCTGA
CAGTTACCAATGCTTAATCAGTGAGGCACCTATCTCAGCG
ATCTGTCTATTTCGTTCATCCATAGTTGCCTGACTCCCCGT
CGTGTAGATAACTACGATACGGGAGGGCTTACCATCTGGC
CCCAGTGCTGCAATGATACCGCGAGACCCACGCTCACCG
GCTCCAGATTTATCAGCAATAAACCAGCCAGCCGGAAGG
GCCGAGCGCAGAAGTGGTCCTGCAACTTTATCCGCCTCC
ATCCAGTCTATTAATTGTTGCCGGGAAGCTAGAGTAAGTA
GTTCGCCAGTTAATAGTTTGCGCAACGTTGTTGCCATTGC
TACAGGCATCGTGGTGTCACGCTCGTCGTTTGGTATGGCT
TCATTCAGCTCCGGTTCCCAACGATCAAGGCGAGTTACAT
GATCCCCCATGTTGTGCAAAAAAGCGGTTAGCTCCTTCGG
TCCTCCGATCGTTGTCAGAAGTAAGTTGGCCGCAGTGTTA
TCACTCATGGTTATGGCAGCACTGCATAATTCTCTTACTGT
CATGCCATCCGTAAGATGCTTTTCTGTGACTGGTGAGTAC
TCAACCAAGTCATTCTGAGAATAGTGTATGCGGCGACCG
AGTTGCTCTTGCCCGGCGTCAATACGGGATAATACCGCGC
CACATAGCAGAACTTTAAAAGTGCTCATCATTGGAAAAC
GTTCTTCGGGGCGAAAACTCTCAAGGATCTTACCGCTGTT
GAGATCCAGTTCGATGTAACCCACTCGTGCACCCAACTG
ATCTTCAGCATCTTTTACTTTCACCAGCGTTTCTGGGTGA
GCAAAAACAGGAAGGCAAAATGCCGCAAAAAAGGGAAT
AAGGGCGACACGGAAATGTTGAATACTCATACTCTTCCTT
TTTCAATATTATTGAAGCATTTATCAGGGTTATTGTCTCATG
AGCGGATACATATTTGAATGTATTTAGAAAAATAAACAAA
TAGGGGTTCCGCGCACATTTCCCCGAAAAGT
